# Intramolecular carbon isotope signals reflect metabolite allocation in plants

**DOI:** 10.1101/2021.06.25.449710

**Authors:** Thomas Wieloch, Thomas David Sharkey, Roland Anton Werner, Jürgen Schleucher

## Abstract

Stable isotopes at natural abundance are key tools to study physiological processes occurring outside the temporal scope of manipulation and monitoring experiments. Whole-molecule carbon isotope ratios (^13^C/^12^C) enable assessments of plant carbon uptake yet conceal information about carbon allocation. Here, we identify an intramolecular ^13^C/^12^C signal at treering glucose C-5 and C-6 and develop experimentally testable theories on its origin. More specifically, we assess the potential of processes within C3 metabolism for signal introduction based *(inter alia)* on constraints on signal propagation posed by metabolic networks. We propose that the intramolecular signal reports carbon allocation into major metabolic pathways in actively photosynthesising leaf cells including the anaplerotic, shikimate, and non-mevalonate pathway. We support our theoretical framework by linking it to previously reported whole-molecule ^13^C/^12^C increases in cellulose of ozone-treated *Betula pendula* and a highly significant relationship between the intramolecular signal and tropospheric ozone concentration. Our theory postulates a pronounced preference of leaf-cytosolic triose-phosphate isomerase to catalyse the forward reaction *in vivo* (dihydroxyacetone phosphate to glyceraldehyde 3-phosphate). In conclusion, intramolecular ^13^C/^12^C analysis resolves information about carbon uptake and allocation enabling more comprehensive assessments of carbon metabolism than whole-molecule ^13^C/^12^C analysis.

**Highlight:** Intramolecular ^13^C/^12^C analysis resolves information about carbon uptake and allocation (and associated environmental controls) enabling more comprehensive assessments of carbon metabolism, plant-environment interactions, and environmental variability than whole-molecule ^13^C/^12^C analysis.

## Introduction

Plant carbon metabolism is a central component of the global carbon cycle. It both depends on and affects environmental properties. Improved understanding of long-term plant-environment interactions relies on information from plant archives (such as tree rings) because manipulation and monitoring experiments can cover short to medium timescales only. Stable carbon isotope (^13^C/^12^C) analysis is among the most advanced tools to extract physiological and environmental information from plant archives. Conventionally, average ^13^C/^12^C ratios of whole plant metabolites are analysed. However, this approach neglects ^13^C/^12^C differences known to occur among individual carbon positions of plant metabolites (Abelson and Hoering, 1961). By contrast, we recently analysed intramolecular ^13^C/^12^C ratios in glucose extracted across an annually-resolved *Pinus nigra* tree-ring timeseries (1961-1995) and reported intramolecular ^13^C signals (i.e., systematic ^13^C/^12^C variation confined to individual glucose carbon positions; Wieloch *et al*., 2018). Only after their ecophysiological origins have been elucidated, may these archived signals become useful for applications within the plant and Earth sciences.

Based on our previous dataset (Wieloch *et al*., 2018), we have already pinpointed a ^13^C signal at tree-ring glucose C-4 and proposed it informs about carbon flux around leaf-cytosolic glyceraldehyde-3-phosphate dehydrogenases and associated energy metabolism (Wieloch, 2021; Wieloch *et al*., 2021). Here, we utilise the same dataset to isolate a ^13^C signal at tree-ring glucose C-5 and C-6. Since intramolecular ^13^C variation is governed *(inter alia)* by enzyme isotope effects and metabolite partitioning (Hayes, 2001), we hypothesise the signal can be linked to shifts in carbon allocation and underlying environmental controls. Thus, we develop experimentally testable theories on ecophysiological mechanisms that can introduce the signal at glucose C-5 and C-6. To this end, we consider all enzyme reactions within central carbon metabolism of C3 plants. This includes the Calvin-Benson cycle (CBC), the photosynthetic carbon oxidation (PCO) cycle, starch and sucrose synthesis and degradation, cellulose synthesis, the pentose phosphate pathway, glycolysis, and carbon metabolism downstream of phospho*enol*pyruvate (PEP). Carbon exchange between other biochemical pathways and the pathway leading to the formation of tree-ring glucose are presumably small, particularly when integrated over the course of growing seasons, the timeframe of tree-ring formation. Thus, these processes cannot introduce ^13^C signals of substantial size into tree-ring archives. Furthermore, we only consider primary isotope effects (which occur at atoms with altered binding after chemical reactions). Sizes of secondary isotope effects (which occur at atoms with unaltered binding after chemical reactions due to indirect involvement in reaction mechanisms) are usually small and therefore unlikely to introduce detectable ^13^C signals into tree-ring archives. Finally, we present evidence supporting our theory. For this part, we reanalyse our own tree-ring dataset (Wieloch et al., 2018) in combination with publicly accessible climate data and ^13^C/^12^C data from an ozone-treatment experiment published by Saurer et al. (1995).

We distinguish two major types of ^13^C fractionation; diffusion-Rubisco fractionation, and post-Rubisco fractionation (Wieloch *et al*., 2018). Diffusion-Rubisco fractionation accompanies CO_2_ diffusion from ambient air into plant chloroplasts and subsequent carbon fixation by Rubisco (Figs. 1–2; Farquhar *et al*., 1982). It affects all carbon positions of plant glucose equally (Wieloch *et al*., 2018). By contrast, post-Rubisco fractionation results from metabolic processes downstream of Rubisco and is position-specific (Figs. 1–3). Deconvolution of the two fractionation types requires the intramolecular approach.

**Figure 1.**
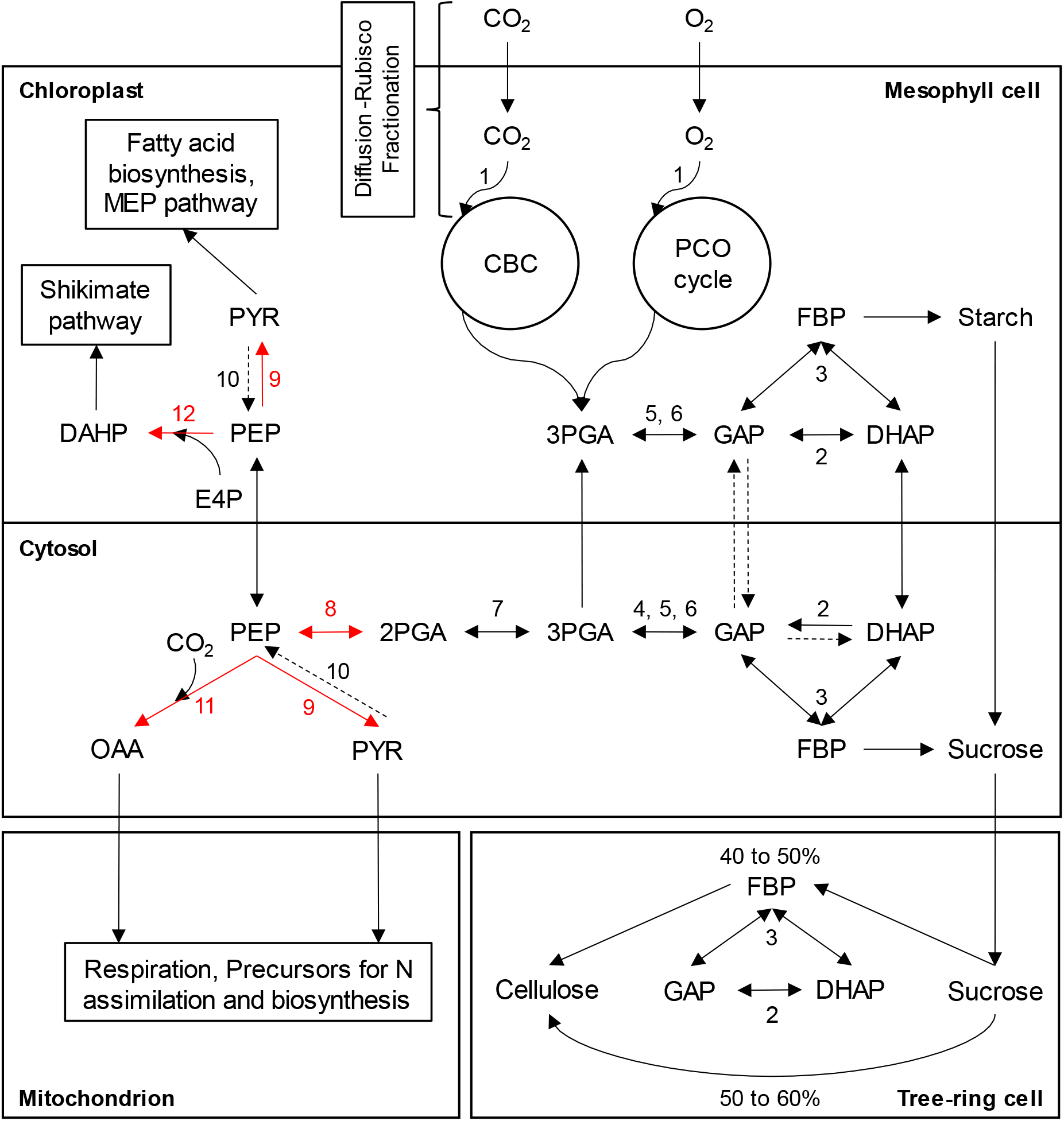
Central carbon metabolism in trees. Solid and dashed arrows represent substantial and negligible metabolite fluxes, respectively. Red arrows: Reactions introducing the *Δ*_5-6_′ signal. Abbreviations: CBC, Calvin-Benson cycle; PCO cycle, photosynthetic carbon oxidation cycle (photorespiration); MEP, non-mevalonate pathway. Metabolites: 2PGA, 2-phosphoglycerate; 3PGA, 3-phosphoglycerate; DAHP, 3-deoxy-D-*arabino*-heptulosonic acid 7-phosphate; DHAP, dihydroxyacetone phosphate; E4P, erythrose 4-phosphate; FBP, fructose 1,6-bisphosphate; GAP, glyceraldehyde 3-phosphate; OAA, oxaloacetate; PEP, phospho*ewo/*pyruvate; PYR, pyruvate. Enzymes: (1) Rubisco, ribulose-1,5-bisphosphate carboxylase/oxygenase; (2) TPI, triose-phosphate isomerase; (3) Aldolase, fructose-bisphosphate aldolase; (4) np-GAPDH, irreversible non-phosphorylating glyceraldehyde-3-phosphate dehydrogenase; (5) p-GAPDH, reversible phosphorylating glyceraldehyde-3-phosphate dehydrogenase; (6) PGK, phosphoglycerate kinase; (7) PGM, phosphoglycerate mutase; (8) enolase, (9) PK, pyruvate kinase; (10) PPDK, pyruvate orthophosphate dikinase; (11) PEPC, phospho*enol*pyruvate carboxylase; (12) DAHPS, *3-Deoxy-D-arabino-* heptulosonate 7-phosphate synthase. Figures 2, and 3 show the reactions in more detail. In contrast to its representation here, parts of the PCO cycle reside outside chloroplasts, in peroxisomes, and mitochondria. Localisation of parts of the shikimate pathway in the cytosol is being debated (Maeda and Dudareva, 2012). To avoid clutter, not all metabolic intermediates are shown. For instance, conversion of 3PGA to GAP proceeds via 1,3-bisphosphoglycerate.

Our work makes several conceptual advances. (i) We show how constraints on signal propagation posed by metabolic networks can be used to narrow down signal origins. (ii) A conceptual model describes how the signal propagates from its origin to other glucose carbon positions and metabolite pools. Due to space restrictions, this model is presented in SI 1. (iii) We revise current theory on plant isotope fractionation by ozone exposure. (iv) The present paper and a companion paper on the C-4 signal (Wieloch *et al*., 2021) develop theories that consider all relevant parts of metabolism and link intramolecular ^13^C signals with specific shifts in carbon allocation and their environmental causes. For isotope signals generated within complex metabolic networks, such comprehensive theories are required as starting point for subsequent tailored experimental tests.

## Material and methods

Intramolecular ^13^C/^12^C ratios in tree-ring glucose of *Pinus nigra* from Vienna (Austria) were reported in Wieloch *et al.* (2018). They are expressed in terms of intramolecular ^13^C discrimination, *Δ_i_*′, where *i* denotes individual carbon positions in tree-ring glucose (Wieloch *et al*., 2018; Abbreviations and symbols in Table 1). In this notation, positive values denote discrimination against ^13^C. The prime denotes measurements subjected to a procedure that removes the ^13^C redistribution effect by triose phosphate cycling (SI 2; Wieloch *et al*., 2018). This correction restores leaf-level ^13^C signals. The dataset comprises six annually resolved timeseries (one per glucose carbon) each covering the period 1961 to 1995 and containing 31 timepoints (*n*=6*31=186).

**Table 1.** Abbreviations, terminology, and identifiers.

| Abbreviation | Metabolite | Identifier |
| --- | --- | --- |
| 1,3BPGA | 1,3-Bisphosphoglycerate | CAS 1981-49-3 |
| 2PGA | 2-Phosphoglycerate | CAS 2553-59-5 |
| 3PGA | 3-Phosphoglycerate | CAS 820-11-1 |
| ADP | Adenosine diphosphate | CAS 58-64-0 |
| ATP | Adenosine triphosphate | CAS 56-65-5 |
| DAHP | 3-Deoxy-D-*arabino*-heptulosonic acid 7-phosphate | CAS 2627-73-8 |
| DHAP | Dihydroxyacetone phosphate | CAS 57-04-5 |
| E4P | Erythrose 4-phosphate | CAS 585-18-2 |
| FBP | Fructose 1,6-bisphosphate | CAS 488-69-7 |
| GAP | Glyceraldehyde 3-phosphate | CAS 591-59-3 |
| NAD<sup>+</sup>, NADH | Nicotinamide adenine dinucleotide | CAS 53-84-9 |
| OAA | Oxaloacetate | CAS 328-42-7 |
| PEP | Phospho*enol*pyruvate | CAS 138-08-9 |
| P<sub>i</sub> | Inorganic phosphate | CAS 14265-44-2 |
| PYR | Pyruvate | CAS 127-17-3 |
| Abbreviation | Enzyme (complex) | Identifier |
|  | Enolase | EC 4.2.1.11 |
| Aldolase | Fructose-bisphosphate aldolase | EC 4.1.2.13 |
| DAHPS | 3-Deoxy-D-*arabino*-heptulosonate 7-phosphate synthase | EC 2.5.1.54 |
| np-GAPDH | Non-phosphorylating glyceraldehyde-3-phosphate dehydrogenase | EC 1.2.1.9 |
| p-GAPDH | Phosphorylating glyceraldehyde-3-phosphate dehydrogenase | EC 1.2.1.12/13 |
| PEPC | Phospho*enol*pyruvate carboxylase | EC 4.1.1.31 |
| PEPCK | Phospho*enol*pyruvate carboxykinase | EC 4.1.1.49 |
| PGK | Phosphoglycerate kinase | EC 2.7.2.3 |
| PGM | Phosphoglycerate mutase | EC 5.4.2.11 |
| PK | Pyruvate kinase | EC 2.7.1.40 |
| PPDK | Pyruvate orthophosphate dikinase | EC 2.7.9.1 |
| Rubisco | Ribulose-1,5-bisphosphate carboxylase/oxygenase | EC 4.1.1.39 |
| TPI | Triose-phosphate isomerase | EC 5.3.1.1 |
| Abbreviation | Other |  |
| CBC | Calvin-Benson cycle |  |
| MEP pathway | Non-mevalonate pathway |  |
| [O<sub>3</sub>] | Tropospheric ozone concentration |  |
| PCO cycle | Photosynthetic carbon oxidation cycle |  |
| *rH* | Relative humidity |  |
| *SD* | Sunshine duration |  |
| TPC | Triose phosphate cycling |  |
| *VPD* | Air vapour pressure deficit |  |
| $\Delta C_i$ | Difference of intercellular CO<sub>2</sub> concentrations | |
| $\delta^{13}C_a$ | <sup>13</sup>C abundance of atmospheric CO<sub>2</sub> | |
| $\Delta\delta^{13}C_p$ | Difference of whole-molecule <sup>13</sup>C abundances of plant matter | |
| $\Delta$ | Whole-molecule <sup>13</sup>C discrimination | |
| $\Delta_i$ | Intramolecular <sup>13</sup>C discrimination | |
| $\Delta_i'$ | Intramolecular <sup>13</sup>C discrimination corrected for TPC | |
| $\Delta\Delta$ | Difference of <sup>13</sup>C discrimination | |
| $\Delta\Delta_{DR}$ | Difference of <sup>13</sup>C discrimination due to diffusion-Rubisco fractionation | |

Additionally, we reanalysed published differences in intercellular CO_2_ concentration, Δ*C*_i_, and whole-molecule ^13^C abundance, Δ*δ*^13^C_p_, between ozone-treated and control plants of *Betula pendula* grown in 1992 (Table 2; 90/40 nl O_3_ 1^-1^ day/night vs. <3 nl O_3_ 1^-1^; Saurer *et a*., 1995). These authors used two different methods to determine Δ*C*_i_. Here, we calculated Δ*C*_i_ averages. Corresponding differences in ^13^C discrimination by the diffusion-Rubisco interface were estimated as

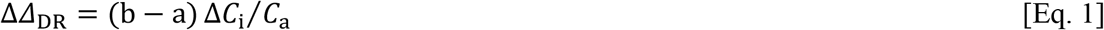

where a and b denote discrimination factors of CO_2_ diffusion (4.4‰) and Rubisco carboxylation (29‰), respectively, and *C*_a_ denotes atmospheric CO_2_ concentration (356 ppm in 1992; Farquhar *et al.*, 1982). Reported Δ*δ*^13^C_p_ values (Table 2) were transformed to Δ*Δ* values as

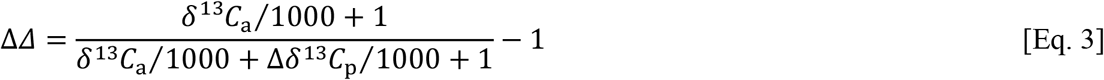

where *R*_a_ denotes the ^13^C/^12^C ratio of atmospheric CO_2_, and Δ*R*_p_ denotes the difference in ^13^C/^12^C ratios between ozone-treated and control plants. To utilise Δ*δ*^13^C_p_ values reported in per mill, we transformed equation 2 as

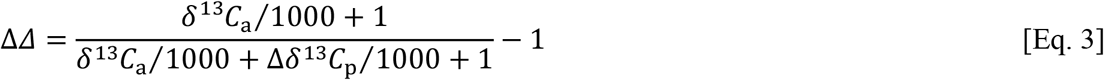

where *δ*^13^C_a_ denotes the ^13^C abundance of atmospheric CO_2_ (−8‰ in 1992; Leuenberger, 2007).

**Table 2.** Reanalysed data originally published by Saurer *et al.* (1995).

|  | C/LF | O<sub>3</sub>/LF | C/HF | O<sub>3</sub>/HF |
| --- | --- | --- | --- | --- |
| $\beta \pm SD$ | $0.006 \pm 0.002$ | $0.013 \pm 0.007$ *** | $0.014 \pm 0.005$ | $0.017 \pm 0.005$ * |
| $\Delta C_i \pm SD$ [ppm] | O<sub>3</sub>/LF - C/LF | | O<sub>3</sub>/HF - C/HF | |
| Steady state | $10 \pm 16$ * | | $34 \pm 41$ ** | |
| Diurnal course | $5 \pm 28$ | | $9 \pm 35$ * | |
| $\Delta \delta^{13}\text{C}_p \pm SD$ [‰] | O<sub>3</sub>/LF - C/LF | | O<sub>3</sub>/HF - C/HF | |
| Leaf cellulose | $1.1 \pm 0.7$ * | | $0.4 \pm 0.9$ | |
| Stem cellulose | $1.3 \pm 0.6$ ** | | $1.1 \pm 0.6$ ** | |

Treatments: control group, C (<3 nl O<sub>3</sub> l<sup>-1</sup>); ozone-treated group, O<sub>3</sub> (90/40 nl O<sub>3</sub> l<sup>-1</sup> day/night); low fertilisation, LF; high fertilisation, HF. $\beta$ denotes the carboxylation rate of phosphoenolpyruvate carboxylase relative to the total carboxylation rate of phosphoenolpyruvate carboxylase and Rubisco measured *in vitro*. $\Delta C_i$ denotes differences in leaf intercellular CO<sub>2</sub> concentrations between ozone-treated and control plants measured by two different methods ('steady state', 'diurnal course'). $\Delta \delta^{13}\text{C}_p$ denotes differences in carbon isotope ratios between ozone-treated and control plants in leaf and stem cellulose. Significance levels of differences between ozone-treated and control plants: \*, $p \leq 0.05$ ; \*\*, $p \leq 0.01$ ; \*\*\*, $p \leq 0.001$ .

For the regression analysis and linear modelling, we used ^13^C/^12^C data from Wieloch *et al.* (2018) and publicly accessible climate data. Data of sunshine duration, *SD,* and relative humidity, *rH*, were acquired from the climate station Hohe Warte in Vienna (Klein Tank *et al*., 2002). Tropospheric ozone concentrations, [*O_3_*], were acquired from Stephansplatz, Laaer Berg, Hermannskogel, Hohe Warte, and Lobau (City of Vienna, Municipal Department 22).

## Results and discussion

### Tree-ring glucose exhibits a post-Rubisco signal at C-5 and C-6

Figure 4 shows results of a Hierarchical Cluster Analysis which groups *Δ_i_*′ timeseries according to co-variability (Wieloch *et al*., 2018). That is, *Δ_i_*′ timeseries carrying common ^13^C signals form clusters. Primary separation occurs between the *Δ*_1_′ to *Δ*_3_′ cluster and the *Δ*_4_’ to *Δ*_6_ cluster. Average timeseries pertaining to these clusters are entirely uncorrelated (*r*=0.08, *p*>0.65, *n*=31).

Thus, these clusters convey entirely different ecophysiological information. Wieloch *et al.* (2018) justified the use of air vapour pressure deficits, *VPD,* as proxy of diffusion-Rubisco fractionation. While the average timeseries of the *Δ_1_* to *Δ_3_* cluster correlates highly significantly with *VPD* (*r*=-0.70, *p*=0.00001, *n*=31), the average timeseries of the *Δ_4_* to *Δ_6_* cluster is not significantly correlated (*r*=-0.30, *p*>0.05, *n*=31). This indicates that the diffusion-Rubisco signal is preserved at glucose C-1 to C-3 but not at C-4 to C-6. Among all *Δ_i_*′, *Δ_5_* and *Δ_6_* exhibit the most significant correlation (*r*=0.61, *p*≤0.001, *n*=31). Since the diffusion-Rubisco signal is confined to glucose C-1 to C-3, we argue C-5 and C-6 exhibit a strong post-Rubisco signal denoted *Δ*_5-6_’ signal.

### Exclusion of metabolic locations as origin of the *Δ*_5-6_′ signal

Much is known about plant carbon metabolism. Based on this knowledge, we can exclude several metabolic locations as origin of the *Δ*_5-6_′ signal as a first step in the theory development. Note that the *Δ*_5-6_′ signal is introduced at the level of three-carbon compounds because reactions at other levels do not modify carbon bonds that become glucose C-5 and C-6.

#### Exclusion of the tree-ring cell as origin of the Δ_5-6_′ signal

^13^C labelling experiments provide compelling evidence for a complete or near complete equilibration of triose phosphates in tree-ring cells of *Quercus robur* (Figs. 1–2; Hill *et al*., 1995). Numerous similar reports for other species and tissues suggest that triose phosphates in the cytosol of heterotrophic tissues are generally substantially equilibrated (Brown and Neish, 1954; Edelman *et al*., 1955; Neish, 1955, 1958; Shafizadeh and Wolfrom, 1955; Seegmiller *et al*., 1956; Altermatt and Neish, 1956; Shibko and Edelman, 1957; McConnell *et al*., 1958; Wolfrom *et al*., 1959; Keeling *et al*., 1988; Hatzfeld and Stitt, 1990; Viola *et al*., 1991). Only two ^13^C labelling studies report no evidence of substantial heterotrophic triose phosphate equilibration (Greathouse, 1953; Kikuta and Erickson, 1969). However, these authors analysed tissues during stages of exceptionally rapid fruit development with either high hexose phosphate flux into cotton-boll cellulose or high triose phosphate flux into avocado lipids (Greathouse, 1953; Brown and Neish, 1954; Kikuta and Erickson, 1969). Taken together, these reports suggest that heterotrophic triose phosphates are generally substantially equilibrated during normal growth.

**Figure 2.**
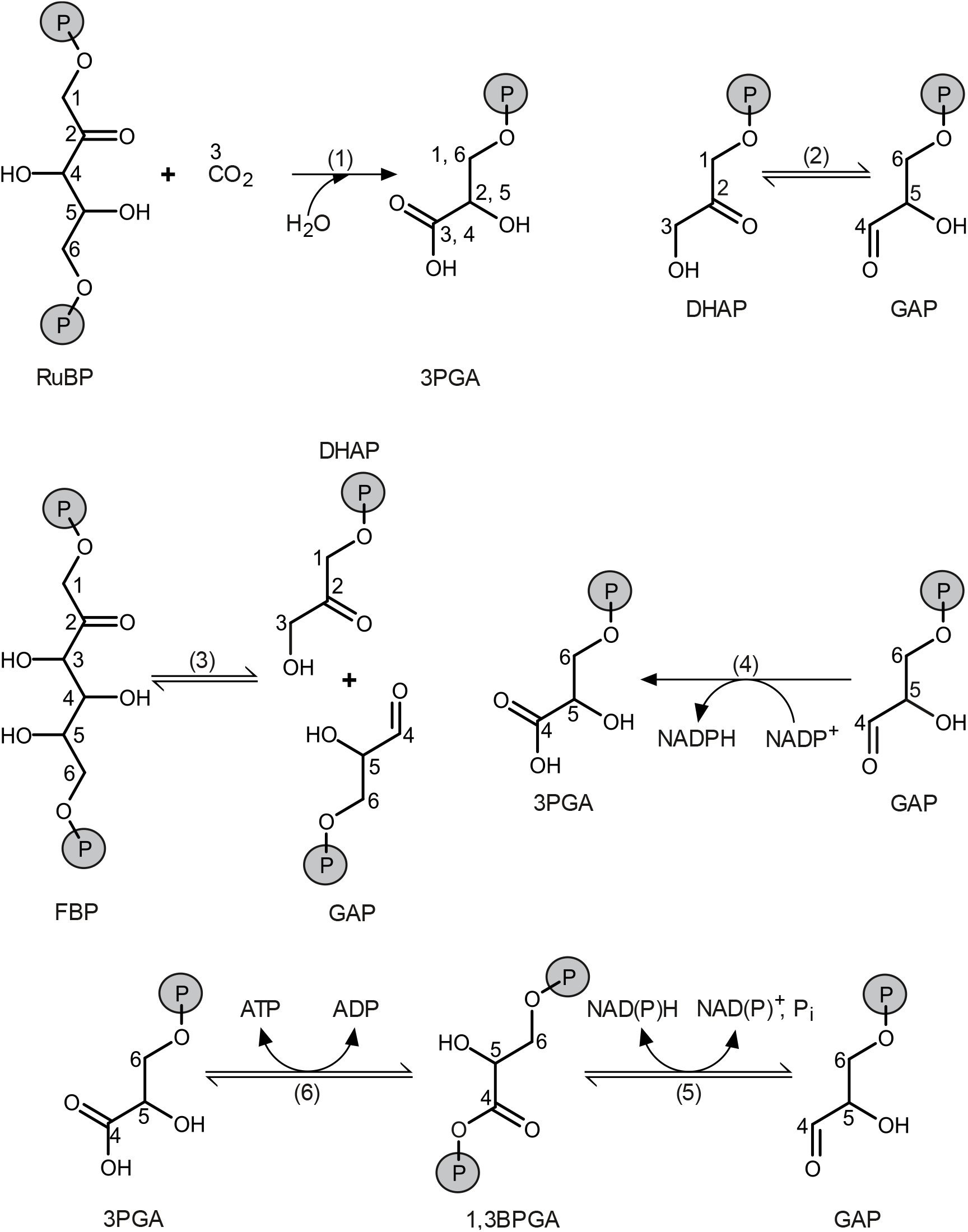
Formation and cleavage of bonds involving carbon atoms in central carbon metabolism. Carbon numbering according to carbon positions in tree-ring glucose. Metabolites: 1,3BPGA, 1,3-bisphosphoglycerate; 3PGA, 3-phosphoglycerate; ADP, adenosine diphosphate; ATP, adenosine triphosphate; DHAP, dihydroxyacetone phosphate; FBP, fructose 1,6-bisphosphate; GAP, glyceraldehyde 3-phosphate; NAD^+^, NADH, nicotinamide adenine dinucleotide; NADP^+^, NADPH, nicotinamide adenine dinucleotide phosphate; P_i_, inorganic phosphate; RuBP, ribulose 1,5-bisphosphate. Numbers in brackets denote enzymes: (1) Rubisco, ribulose-1,5-bisphosphate carboxylase/oxygenase; (2) TPI, triose-phosphate isomerase; (3) Aldolase, fructose-bisphosphate aldolase; (4) np-GAPDH, irreversible nonphosphorylating glyceraldehyde-3-phosphate dehydrogenase; (5) p-GAPDH, reversible phosphorylating glyceraldehyde-3-phosphate dehydrogenase; (6) PGK, phosphoglycerate kinase.

The raw dataset of intramolecular ^13^C discrimination, *Δ_i_*′, in tree-ring glucose (Wieloch *et al*., 2018) exhibits significant correlations among all pairs of symmetry-related timeseries (Table 3). That is, significant correlations occur between *Δ*_1_ and *Δ*_6_, *Δ*_2_ and *Δ*_5_, and *Δ*_3_ and *Δ*_4_. These correlations likely result from carbon redistribution by triose phosphate cycling (TPC) which involves triose phosphate equilibration. Wieloch *et al.* (2018) describe this process mathematically and used the model to remove the TPC effect from *Δ_i_*′ yielding a TPC-free dataset, *Δ_i_*′. In this latter dataset, significant correlations among pairs of symmetry-related timeseries are absent (Table 4). This provides strong evidence for the occurrence of substantial triose phosphate equilibration in tree-ring cells of the samples discussed here.

**Table 3.**
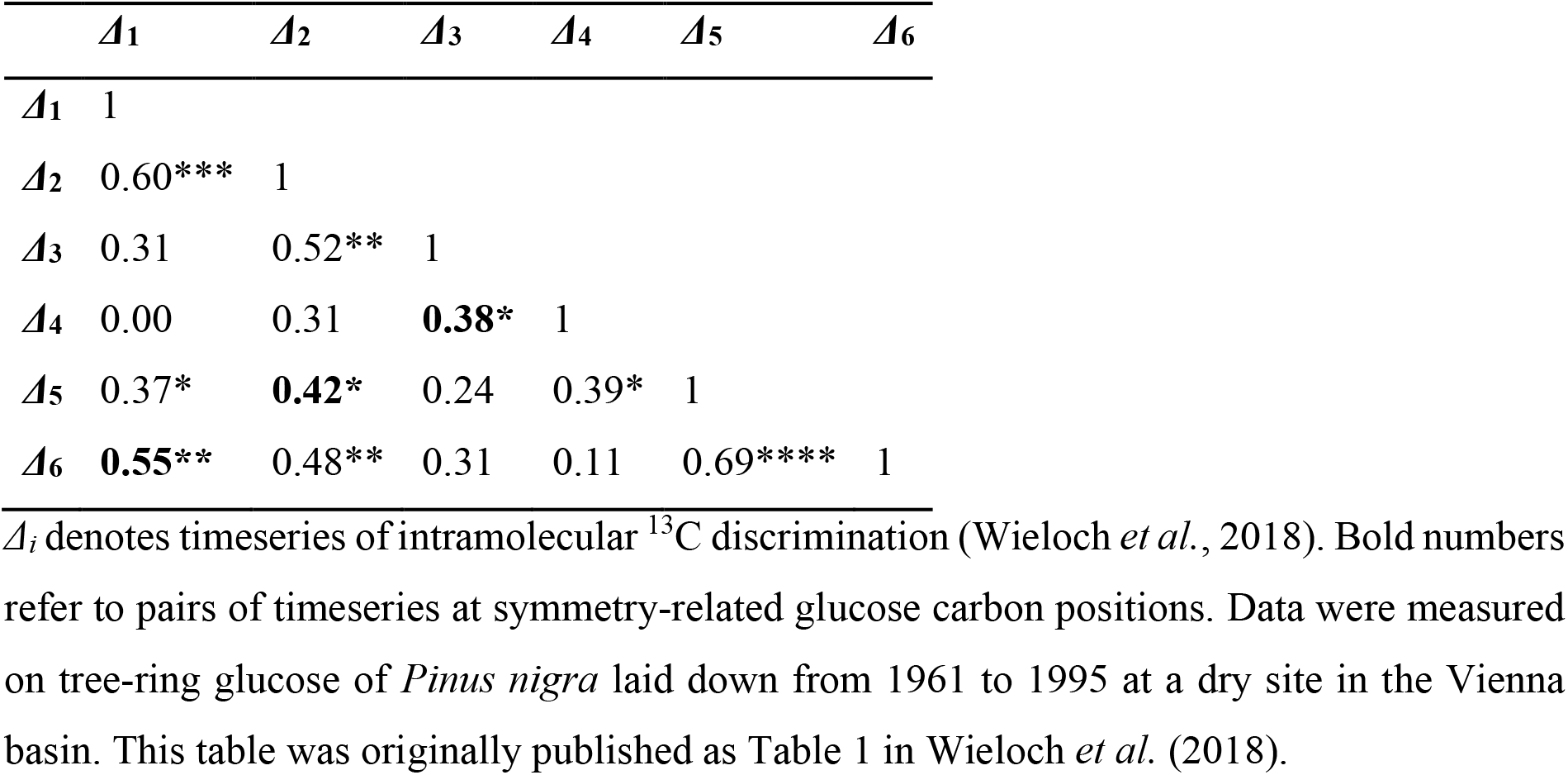
Correlation coefficients and significance levels obtained from cross-correlation analysis on *Δi* (*,*p*≤0.05; **,*p*≤0.01; ***,*p*≤0.001; ****,*p*≤0.0001; *n*=6*31).

**Table 4.** Correlation coefficients and significance levels obtained from cross-correlation analysis on *Δ_i_*′ (*,*p*≤0.05; **,*p*≤0.01; ***,*p*≤0.001; *n*=6*31).

| | $\Delta_1'$ | $\Delta_2'$ | $\Delta_3'$ | $\Delta_4'$ | $\Delta_5'$ | $\Delta_6'$ |
| --- | --- | --- | --- | --- | --- | --- |
| $\Delta_1'$ | 1 | | | | | |
| $\Delta_2'$ | 0.54** | 1 | | | | |
| $\Delta_3'$ | 0.31 | 0.48** | 1 | | | |
| $\Delta_4'$ | -0.12 | 0.10 | **-0.12** | 1 | | |
| $\Delta_5'$ | 0.11 | **-0.07** | 0.03 | 0.32 | 1 | |
| $\Delta_6'$ | **0.08** | 0.19 | 0.21 | 0.06 | 0.61*** | 1 |

If a process in tree-ring cells had introduced a signal at carbon positions corresponding to glucose C-5 and C-6, triose phosphate equilibration would have transmitted it to carbon positions corresponding to glucose C-2 and C-1. The signal at C-5 would have had the same size as the signal at C-2 and the signal at C-6 would have had the same size as the signal at C-1. Please note that equally-sized signals at symmetry-related glucose carbon positions are not removed by the method removing TPC effects (Wieloch *et al*., 2018). Since the *Δ*_5-6_′ signal is absent in *Δ*_1_ and *Δ*_2_′ (Fig. 4), it must have been introduced at the leaf level.

#### Exclusion of the CBC and PCO cycle as origin of the Δ_5-6_′ signal

Introduction of the *Δ*_5-6_′ signal within the CBC or PCO cycle can be excluded because hexose phosphate synthesis includes conversion of photosynthetic/photorespiratory glyceraldehyde 3-phosphate (GAP, *Δ*_4_ to *Δ4)* to dihydroxyacetone phosphate (DHAP, *Δ*_3_ to *Δ*_1_′) by triosephosphate isomerase (TPI, Figs. 1–2). This would transmit any ^13^C signal present at GAP carbon positions corresponding to glucose C-5 and C-6 to DHAP carbon positions corresponding to glucose C-2 and C-1. More generally, metabolites feeding into the stromal GAP pool can be excluded as origin of the *Δ*_5-6_′ signal based on the same reasoning.

#### Exclusion of reactions downstream of OAA, pyruvate, and DAHP as Δ_5-6_′ signal origin

Pyruvate kinase (PK) and pyruvate orthophosphate dikinase (PPDK) interconvert PEP and pyruvate (Figs. 1, and 3). The PK reaction is strongly on the side of pyruvate and considered nearly irreversible (Nageswara Rao *et al*., 1979; Tcherkez *et al*., 2011). In illuminated leaves of C3 plants, PPDK activity is either very low or undetectable, except for orchids, and grasses (Hocking and Anderson, 1986). In illuminated leaves of *Xanthium strumarium,* flux from pyruvate to PEP is very small at ≈0.05% of net CO_2_ assimilation (Tcherkez *et al*., 2011). In *Arabidopsis thaliana,* PPDK is upregulated during leaf senescence which is believed to facilitate nitrogen remobilisation (Taylor *et al*., 2010). This is of minor importance here because leaf senescence occurs during a short period relative to the multi-year lifespan of conifer needles. In *Nicotiana tabacum,* PPDK activity is increased up to 2.7-fold under strong drought (Doubnerová Hýsková *et al*., 2014). However, this should not result in significant flux in relation to fluxes in carbohydrate metabolism since basal PPDK activities in C3 plants are generally low (Hocking and Anderson, 1986; Tcherkez *et al*., 2011). Thus, flux from pyruvate to PEP should be small, and transmission of ^13^C signals in pyruvate to cytosolic carbohydrates by the PK/PPDK interface should be negligible.

**Figure 3.**
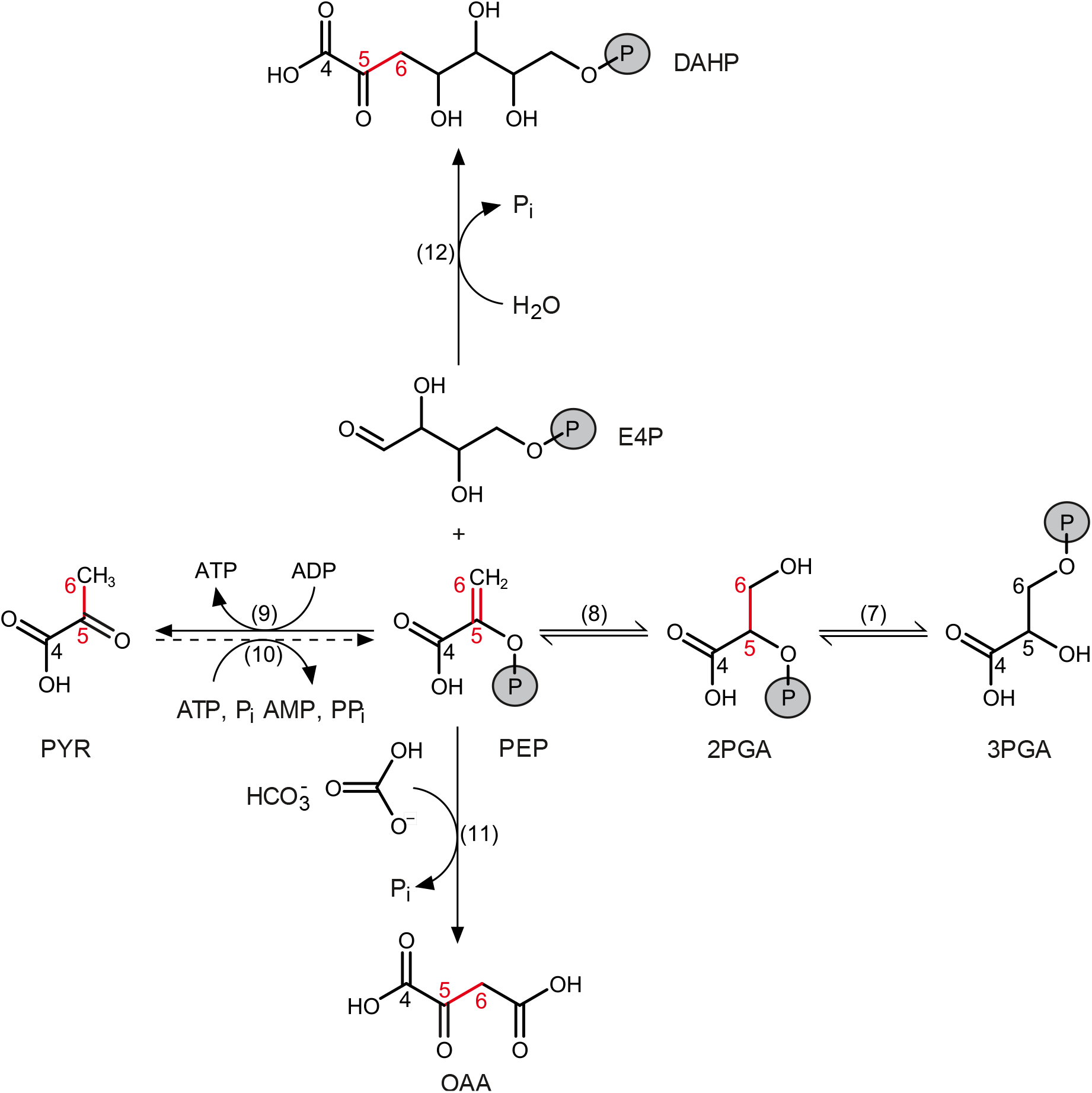
Formation and cleavage of bonds involving carbon atoms in central carbon metabolism. Solid and dashed arrows represent substantial and negligible metabolite fluxes, respectively. Red: Carbon bond modifications possibly accompanied by primary isotope effects. Carbon numbering according to carbon positions in tree-ring glucose. Metabolites: 2PGA, 2-phosphoglycerate; 3PGA, 3-phosphoglycerate; AMP, adenosine monophosphate; ADP, adenosine diphosphate; ATP, adenosine triphosphate; DAHP, 3-deoxy-D-*arabino*-heptulosonic acid 7-phosphate; E4P, erythrose 4-phosphate; 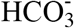, bicarbonate; OAA, oxaloacetate; PEP, phospho*enol*pyruvate; P_i_, inorganic phosphate; PP_i_, pyrophosphate; PYR, pyruvate. Numbers in brackets denote enzymes: (7) PGM, phosphoglycerate mutase; (8) enolase, (9) PK, pyruvate kinase; (10) PPDK, pyruvate orthophosphate dikinase; (11) PEPC, phospho*enol*pyruvate carboxylase; (12) DAHPS, 3-Deoxy-D-*arabino*-heptulosonate 7-phosphate synthase.

Phospho*enol*pyruvate carboxylase (PEPC) and phospho*enol*pyruvate carboxykinase (PEPCK) interconvert PEP and oxaloacetate (OAA). Conversion of PEP to OAA by PEPC is irreversible (Chollet *et al*., 1996). To our knowledge, there are no reports of PEPCK activity in mesophyll cells where bulk carbohydrate synthesis takes place (Pyke, 2001). PEPCK RNA and protein were not detected in leaves of *Solanum lycopersicum* irrespective of their developmental stage (Bahrami *et al*., 2001; Famiani *et al*., 2016). PEPCK protein or activity were not detected in leaves of *Hordeum vu/gare* (Chen *et al*., 2000). In mature leaves of *Arabidopsis thaliana,* PEPCK protein amount and activity are low and likely confined to specific cell types (Malone *et al*., 2007). In leaves of *Nicotiana tabacum,* PEPCK occurs in trichomes and stomata (Leegood *et al*., 1999; Malone *et al*., 2007). In leaves of *Cucumis sativus,* PEPCK occurs in trichomes and phloem cells (Leegood *et al*., 1999; Chen *et al*., 2004). In leaves of *Oryza sativa*, PEPCK occurs in hydathodes, stomata, and the vascular parenchyma (Bailey and Leegood, 2016). Thus, transmission of ^13^C signals in OAA to cytosolic carbohydrates by the PEPC/PEPCK interface should not occur.

3-Deoxy-D-*arabino*-heptulosonate 7-phosphate synthase (DAHPS) catalyses the irreversible reaction from PEP and erythrose 4-phosphate (E4P) to 3-deoxy-D-*arabino*-heptulosonic acid 7-phosphate (DAHP; Herrmann, 1995). To our knowledge, there are no reports of enzymes catalysing the back reaction. Thus, transmission of ^13^C signals in DAHP to cytosolic carbohydrates should not occur.

Taken together, leaf-level carbon fluxes from OAA, pyruvate, and DAHP to PEP are negligible or absent (Figs. 1, and 3). Therefore, reactions downstream of OAA, pyruvate, and DAHP cannot feed significant amounts of carbon and associated ^13^C signals into carbohydrate metabolism.

#### Exclusion of starch, sucrose, and cellulose metabolism, and the pentose phosphate pathway as origin of the Δ_5-6_′ signal

Reactions leading directly from stromal GAP to the formation of starch, sucrose, and cellulose, reactions remobilising starch, and reactions of the pentose phosphate pathway do not simultaneously modify carbon bonds that become glucose C-5 and C-6. This excludes these pathways for *Δ*_5-6_′ signal introduction.

### Origin of the *Δ*_5-6_′ signal

After excluding several metabolic locations as the origin of the *Δ*_5-6_′ signal (see ‘Exclusion of metabolic locations as origin of the *Δ*_5-6_′ signal’), the glycolytic pathway and PEP metabolism in leaves are left for consideration. Within this system, cytosolic GAP is used for PEP metabolism and sucrose synthesis (Fig. 1). Thus, GAP constitutes a central branch point in leaf carbon metabolism enabling isotope fractionation.

#### Leaf-level enolase, PEPC, PK, and/or DAHPS introduce the Δ_5-6_′ signal

Enolase, PEPC, PK, and DAHPS are the only enzymes which simultaneously modify carbon bonds that become glucose C-5 and C-6 (Figs. 1–3). Reactions catalysed by these enzymes may be accompanied by ^13^C effects of substantial size and may thus introduce the *Δ*_5-6_′ signal. Enolase interconverts 2-phosphoglycerate (2PGA) and PEP (Figs. 1, and 3). *In vivo,* the reaction operates close to equilibrium (Kubota and Ashihara, 1990) and might thus be accompanied by an equilibrium isotope effect. Formation of the C=C double bond in PEP likely favours turnover of ^13^C isotopologues of 2PGA leading to ^13^C enrichment in PEP. A ^13^C signal might then arise from varying allocation of PEP to downstream processes (Fig. 1). Increased downstream consumption would remove more ^13^C-enriched PEP and leave behind more ^12^C-enriched 2PGA for glucose synthesis.

Kinetic isotope effects may accompany the unidirectional conversions of PEP to OAA by PEPC, PEP to pyruvate by PK, and PEP and E4P to DAHP by DAHPS (Figs. 1, and 3). These reactions break the C=C double bond in PEP and can therefore be expected to favour turnover of ^12^C-isotopologues of PEP leaving behind ^13^C-enriched PEP for glucose synthesis. Due to the usually larger size of kinetic isotope effects compared to equilibrium isotope effects, effects by PEPC, PK and DAHPS can be expected to outweigh any reciprocal effect by enolase. Thus, considering all four enzymes together, increasing turnover of PEP by PEPC, PK, and DAHPS can be expected to result in ^13^C-enriched tree-ring glucose, i.e., *Δ*_5-6_′ decreases.

#### Signal transmission to tree-ring glucose

Isotope signals generated at the level of leaf-cytosolic PEP or 2PGA need to be transmitted to GAP to then enter hexose phosphates and tree-ring glucose (Fig. 1). Transmission of a signal introduced by cytosolic enzymes is straightforward since the cytosolic glycolytic reactions between PEP and GAP are at equilibrium (Kubota and Ashihara, 1990). Transmission of a signal introduced by stromal enzymes is more intricate. First, it requires an incomplete or low-activity glycolytic pathway in leaf chloroplasts because signal equilibration with stromal triose phosphates would result in even signal distribution over all glucose carbon positions (SI 1.6). An incomplete glycolytic pathway in leaf chloroplasts is supported by a reported lack of enolase in *Arabidopsis thaliana* and *Oryza sativa* (van der Straeten *et al*., 1991; Prabhakar *et al.*, 2009; Fukayama *et al*., 2015). Second, signal transmission from stromal PEP to C-5 and C-6 of cytosolic hexose phosphate requires chloroplast export of PEP. Transport of PEP across the chloroplasts’ inner membrane is mediated by the PEP/Pi translocator as counter-exchange with P_i_, PEP, or 2PGA and the putative *in vivo* preference for the transport of P_i_, and PEP (Fischer *et al*., 1997; Flügge *et al*., 2011). Numerous stromal processes, such as the shikimate pathway and fatty acid biosynthesis, rely on PEP import from the cytosol (Streatfield *et al*., 1999; Flügge *et al*., 2011). Therefore, a net flux of PEP from the cytosol to chloroplasts can be expected. However, members of the phosphate translocator family are believed to be highly inefficient. For instance, merely 10% of the activity of the triose phosphate translocator is used for net export of triose phosphate from chloroplasts; 90% is wasted on futile counter-exchanges (Flügge, 1987, 1999). In addition, low stromal and high cytosolic P_i_ levels (Sharkey and Vanderveer, 1989) can be expected to promote chloroplast export of PEP. Consequently, efficient equilibration of cytosolic and stromal PEP pools, and ^13^C signals therein, can be expected. Thus, both cytosolic and stromal enzymes may contribute to the *Δ*_5-6_′ signal.

#### Signal introduction requires substantial carbon fluxes and flux variability

For the introduction of a ^13^C signal, a substantial share of the photosynthetically fixed carbon must be directed towards enolase, PEPC, PK, and/or DAHPS and their downstream derivatives. This share must vary substantially; in the present case, on the interannual timescale. Therefore, we will now discuss carbon fluxes through enolase, PEPC, PK, and DAHPS.

Commonly, PEPC is localised in the cytosol both in dispersion and bound to the outer mitochondrial membrane (Figs. 1, and 3; O’Leary *et al*., 2011). In leaf mesophyll cells of *Oryza sativa,* a putatively rare additional isoform occurs in chloroplasts (Masumoto *et al*., 2010; O’Leary *et al*., 2011). In C3 plants, PEPC provides OAA to replenish tricarboxylic acid cycle intermediates, support nitrogen assimilation, and biosynthetic processes (O’Leary *et al*., 2011; O’Leary and Plaxton, 2017). On average, leaf carbon fixation by PEPC is believed to account for up to 5% of net CO_2_ assimilation (Melzer and O’Leary, 1987). Up-regulation of PEPC occurs (*inter alia*) with drought, salinity, ozone, nitrogen assimilation, and virus infections (see ‘Ecophysiological effects’; O’Leary *et al*., 2011). For instance, ozone triggers both an upregulation of PEPC and a down-regulation of Rubisco (Dizengremel, 2001). In forest trees, the Rubisco/PEPC activity ratio can change from up to 25 in ozone-free air to about 2 under realistic levels of ambient ozone redirecting carbon flux to maintenance and repair processes (Dizengremel, 2001).

Isoforms of PK are localised in both the cytosol and chloroplasts (Figs. 1, and 3; Ambasht and Kayastha, 2002). They provide pyruvate (*inter alia*) for mitochondrial respiration, fatty acid biosynthesis, and the non-mevalonate pathway (MEP). To our knowledge, estimates of the respiratory flux via PK in actively photosynthesising leaves are unavailable. However, this flux may be substantial when photorespiration is low and thus co-vary with photorespiration and its environmental controls (SI 3).

In illuminated photosynthetic tissue of *Arabidopsis thaliana*, fatty acid biosynthesis can occur at a rate of 2.3 μmol C mg chl^-1^ h^-1^ (Bao *et al.,* 2000). Based on this, we estimate a ≈2% carbon flux relative to net CO_2_ assimilation into fatty acid biosynthesis (SI 4). In leaves, this flux is predominantly controlled at the level of acetyl-CoA carboxylase (Page *et al*., 1994; Harwood, 2005; Ohlrogge *et al*., 2015). It responds to the stromal redox state (energy status) and associated environmental controls (Rawsthorne, 2002; Harwood, 2005; Geigenberger and Fernie, 2014).

The plastid-localised MEP pathway is yet another metabolic route carrying substantial flux. With isoprene as a major pathway product in some trees, it commonly consumes ≈2% of net assimilated CO_2_ (Sharkey and Yeh, 2001). In forest trees, high temperature can enhance this fraction to up to 15% (Sharkey *et al*., 1996); a plant response believed to mitigate short-term high-temperature stress (Sharkey and Yeh, 2001).

DAHPS, located in both chloroplasts and the cytosol, is the first enzyme of the shikimate pathway (Figs. 1, and 3; Maeda and Dudareva, 2012). In vascular plants, 20 to 50% of the photosynthetically fixed carbon enters the pathway (Tohge *et al*., 2013). In trees, most of the flux can be expected to occur in heterotrophic tissues supporting lignin biosynthesis. To our knowledge, flux estimates for actively photosynthesising leaves are unavailable. However, the shikimate pathway provides precursors for (*inter alia*) the aromatic amino acids phenylalanine, tryptophan, tyrosine, and their numerous derivatives. Thus, it should carry substantial flux in most tissues. In leaves of *Prunus persica* fed ^13^CO_2_, <6% of the label accumulated in a metabolite fraction comprising lipids, proteins, and residual compounds (Escobar-Gutiérrez and Gaudillère, 1997). Since the shikimate pathway contributes to the biosynthesis of this metabolite fraction among other pathways, its flux must be markedly below 6% of net assimilated CO_2_. In leaves of *Helianthus annuus,* Abadie *et al.* (2018) reported a flux of ≈1% relative to net CO_2_ assimilation into the shikimate pathway product chlorogenate under normal growing conditions. Regulation of the shikimate pathway is primarily exerted by gene expression and post-translational modification in response to developmental and environmental cues (Entus *et al*., 2002; Mir *et al*., 2015). Relative carbon flux through the shikimate pathway can be expected to *(inter alia)* vary with light (Henstrand *et al*., 1992; Logemann *et al*., 2000; Entus *et al*., 2002), ozone (Janzik *et al*., 2005; Betz *et al*., 2009), physical wounding (Dyer *et al*., 1989; Keith *et al*., 1991), bacterial infection (Keith *et al*., 1991; Truman *et al*., 2006), fungal infestation (McCue and Conn, 1989; Henstrand *et al*., 1992; Görlach *et al*., 1995; Bischoff *et al*., 1996, 2001; Ferrari *et al*., 2007), and nitrogen availability (Scheible *et al*., 2004). For instance, in leaves of *Nicotiana tabacum,* induction of DAHPS increased up to 5-fold under ozone fumigation (160 nl l^-1^), and an increase in flux through the shikimate pathway was corroborated by increased levels of pathway products (Janzik *et al.*, 2005). Performing an 83-days ozone fumigation experiment (160 to 190 nl l^-1^, 8 h day^-1^), Betz *et al.* (2009) reported evidence for increased carbon flux into the shikimate pathway in leaves of *Fagus sylvatica.*

Since PEPC, PK, and DAHPS are located downstream of enolase (Figs. 1, and 3), all four enzymes may contribute to the *Δ*_5-6_′ signal. Based on arguments given above, associated carbon fluxes and their variability can be expected to be substantial. Other leaf-level pathways consuming PEP, such as the cytosolic mevalonate pathway, may exert additional control over the *Δ*_5-6_′ signal.

#### Ecophysiological effects

The *Δ*_5-6_′ signal is independent of the diffusion-Rubisco signal at C-1 and C-2 (Fig. 4). Since diffusion-Rubisco fractionation initially affects all carbon entering glucose synthesis equally (see ‘Introduction’), we propose the *Δ*_5-6_′ signal exhibits two components of variance. The first component is inversely correlated with diffusion-Rubisco fractionation and removes the diffusion-Rubisco signal from glucose C-5 and C-6. The second component constitutes systematic variation independent of diffusion-Rubisco fractionation. In the following, we propose ecophysiological mechanisms for the introduction of each component starting with the independent component. Please note that, in the present case, the *Δ*_5-6_′ signal can be expected to be under environmental rather than developmental control (SI 5).

**Figure 4.**
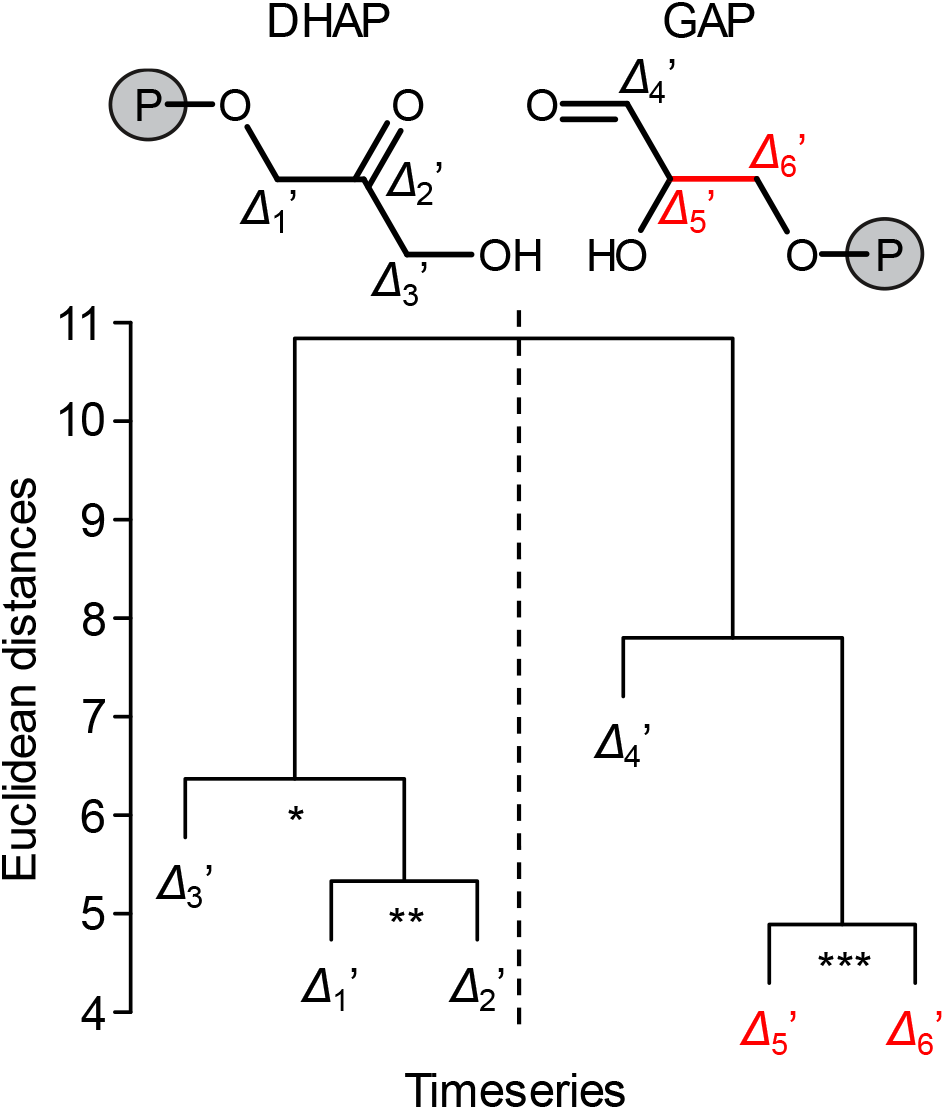
Clustering of *Δ_i_*′ timeseries due to co-variability. *Δ_i_*′ denotes timeseries of intramolecular ^13^C discrimination corrected for triose phosphate cycling (Wieloch *et al*., 2018). Red: Timeseries discussed here. Data were measured on tree-ring glucose of *Pinus nigra* laid down from 1961 to 1995 at a dry site in the Vienna basin (*n*=6*31). Members of clusters marked by asterisks are correlated at the following significance levels: p≤0.05 (*), 0.01 (**), and 0.001 (***). Precursors of tree-ring glucose, dihydroxyacetone phosphate (DHAP) and glyceraldehyde 3-phosphate (GAP) are shown as molecular structures. Modified figure from Wieloch *et al.* (2018).

Wieloch *et al.* (2018) studied effects of *VPD*, precipitation, soil moisture, temperature, and global radiation on the diffusion-Rubisco signal in their *Pinus nigra* samples. These authors found that *VPD*, a measure of environmental drought, exerts predominant control and pointed out that this agrees with expectations for the generally dry study site. Thus, the independent component of the *Δ*_5-6_′ signal is governed by environmental factors other than *VPD.*

The study site is ≈10 km away from the city centre of Vienna and frequently exposed to substantial levels of tropospheric ozone (Oltmans *et al*., 1998; Ainsworth *et al*., 2012). Lefohn (1992) classified *Pinus nigra* as an ozone-sensitive tree species. Radiation stimulates the photochemical reactions of ozone formation (Ainsworth *et al*., 2012). Ozone triggers relative flux increases through the anaplerotic and shikimate pathways via PEPC and DAHPS, respectively (see ‘Signal introduction requires substantial carbon fluxes and flux variability’) and may thus cause ^13^C increases at PEP carbon positions that become glucose C-5 and C-6 (see ‘Leaf-level enolase, PEPC, PK, and/or DAHPS introduce the *Δ*_5-6_′ signal’). This may introduce an isotope signal independent of the diffusion-Rubisco signal due to independence at the level of environmental controls.

By contrast, a process mitigating ozone entry into plant leaves may explain the component of the *Δ*_5-6_′ signal which is inversely correlated with the diffusion-Rubisco signal. Dizengremel (2001) proposed that drought and ozone combined is a main recurring stress factor in forest ecosystems. Isohydric plant species, such as *Pinus nigra*, respond to drought by closing their stomata (Sade *et al*., 2012). Reduced stomatal conductance impedes ozone uptake (Tingey and Hogsett, 1985; Dobson *et al*., 1990; Dizengremel, 2001). In needles of *Pinus halepensis,* PEPC activities in control plants and plants exposed to mild drought stress were similar, strongly increased under ozone stress, but significantly less so under combined ozone and drought stress (Fontaine *et al*., 2003). Thus, anaplerotic flux rates can be expected to be highest under ozone stress but lower when ozone stress is accompanied by drought. While drought causes ^13^C enrichments at all glucose carbon positions due to diffusion-Rubisco fractionation (Wieloch *et al.*, 2018), it can be expected to reduce ozone-induced ^13^C enrichments at glucose C-5 and C-6. This drought component of the ozone response may remove the diffusion-Rubisco signal from glucose C-5 and C-6. In SI 3, we discuss how changes in substrate supply to mitochondrial oxidative phosphorylation (glycolytic pyruvate versus photorespiratory glycine) may additionally contribute to the component of the *Δ*_5-6_′ signal that is inversely correlated with diffusion-Rubisco fractionation.

### Experimental evidence

#### Effects of tropospheric ozone on whole-molecule ^13^C/^12^C composition of plant cellulose

Growing *Betula pendula* at increased ozone levels, several authors reported decreased ^13^C discrimination, *Δ*, in leaf and stem cellulose (Matyssek *et al*., 1992; Saurer *et al*., 1995). Intriguingly, these *Δ* decreases coincided with increased ratios of intercellular to ambient CO_2_ concentrations, *C*_i_/*C*_a_. As pointed out by Matyssek *et al.* (1992) and Saurer *et al.* (1995), this cannot be explained by the standard model of diffusion-Rubisco fractionation which predicts a positive correlation between *C*_i_/*C*_a_ and *Δ* (Farquhar *et al*., 1982). Thus, post-Rubisco fractionation can be expected to cause these ozone-related isotope effects.

Matyssek *et al.* (1992) and Saurer *et al.* (1995) proposed increased relative carbon fixation by PEPC due to ozone explains the *Δ* decreases because carbon fixed by PEPC is strongly ^13^C enriched compared to carbon fixed by Rubisco (Melzer and O’Leary, 1987). While this proposal is in line with significantly increased relative PEPC activities observed under ozone (Table 2), it conflicts with the setup of carbon metabolism. PEPC-fixed carbon supplies downstream metabolism, yet no pathway carrying substantial flux exists that could transfer it into carbohydrate metabolism (see ‘Exclusion of reactions downstream of OAA, pyruvate, and DAHP as *Δ*_5-6_′ signal origin’). Above, we propose an ozone-dependent mechanism for the introduction of the *Δ*_5-6_′ signal which reconciles observations of Matyssek *et al.* (1992) and Saurer *et al.* (1995) with the setup of carbon metabolism (see ‘Ecophysiological effects’).

Saurer *et al.* (1995) reported differences in intercellular CO_2_ concentration, Δ*C*_i_, and whole-molecule ^13^C discrimination, Δ*Δ*, between ozone-treated and control plants. Plants grown with lower amounts of fertiliser (LF) exhibited Δ*C*_i_=7.5 ± 2.6SE ppm, while plants grown with higher amounts of fertiliser (HF) exhibited Δ*C*_i_=21.5 ± 4.5SE ppm. This corresponds to estimated increases in ^13^C discrimination by the diffusion-Rubisco interface of Δ*Δ*_DR_=0.52 ± 0.18SE ‰ and 1.49 ± 0.31SE ‰, respectively (Fig. 5, dashed bars, Eq. 1). However, Saurer *et al.* (1995) reported Δ*Δ* decreases in leaf and stem cellulose under both fertilisation regimes (Fig. 5, solid bars, Eqs. 2–3). With respect to Δ*Δ*_DR_, these decreases are statistically significant (one-tailed t-test: *p*<0.05) except for leaf cellulose synthesised under HF conditions which comes, however, close to being statistically significant (*p*<0.08).

**Figure 5.**
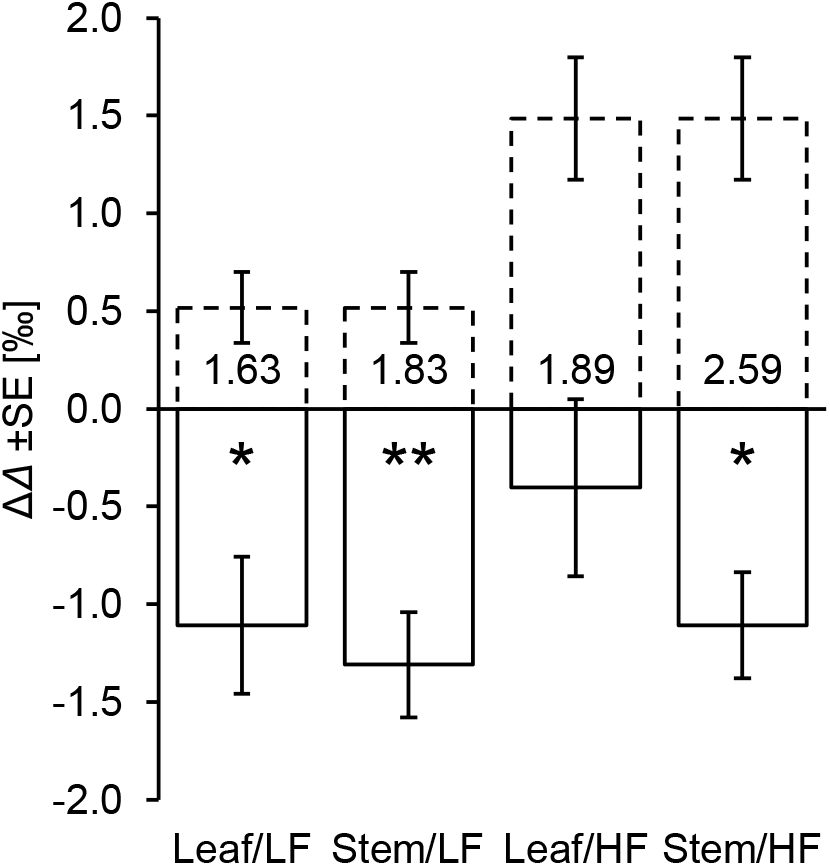
Differences in ^13^C discrimination between ozone-treated and control plants, Δ*Δ* (90/40 day/night versus <3 nl O_3_ 1^-1^). ‘Leaf and ‘Stem’ refers to leaf and stem cellulose of *Betula pendula,* respectively. LF and HF refers to plants grown with low and high amounts of fertiliser, respectively. Solid bars: Measured Δ*Δ* values. Dashed bars: Expected Δ*Δ* values. Expected Δ*Δ* values were estimated using a model by Farquhar *et al.* (1982). This model describes ^13^C discrimination associated with plant carbon uptake including CO_2_ diffusion into plant leaves and assimilation by Rubisco. Numbers inside bars denote differences between measured and expected Δ*Δ* values. Statistically significant differences are marked by asterisks (one-tailed t-test: *, *p*<0.05; **, *p*<0.01). The difference of the Leaf/HF treatment is close to being statistically significant (*p*<0.08). This analysis is based on data published by Saurer *et al.* (1995).

In *Betula pendula*, post-Rubisco fractionation causes average whole-molecule Δ*Δ* decreases of ≈-1.98 ± 0.58SE ‰ (Fig. 5). Below, we propose a fraction of the *Δ*_5-6_′ signal enters glucose C-1 to C-4 through indirect signal propagation via chloroplast metabolism (see ‘Signal propagation to all glucose carbons via chloroplast metabolism’). We estimate the signal at C-5 and C-6 is 6.625-fold larger than at C-1 to C-4 (SI 1.8). Thus, a ≈-1.98 ± 0.58SE ‰ effect at the whole-molecule level scales to ≈-4.56 ± 1.34SE ‰ effects at cellulose glucose C-5 and C-6 and to ≈-0.69 ± 0.20SE ‰ effects at C-1 to C-4 (SI 6). In *Pinus nigra,* measured *Δ*_5-6_′ values fall within a 5.80 ± 1.55SE ‰ range (maximum=22.71 ± 0.99SE ‰, minimum=16.91 ± 0.56SE ‰). Wieloch *et al.* (2018) estimated the *Δ*_5-6_′ timeseries contains 79% systematic and 21% error variance. Assuming the error is fully expressed in both the maximum and minimum value, we estimate a systematic timeseries range of ≈4.58 ± 1.22SE ‰ (5.80 ± 1.55SE ‰ * 0.79). This largely equals the estimated effect at glucose C-5 and C-6 in ozone-treated *Betula pendula* corroborating the theory proposed above. Notably, occurrence of the post-Rubisco fractionation effect in leaf cellulose of *Betula pendula* corroborates the proposed leaf-level origin of the *Δ*_5-6_′ signal.

#### Effect of tropospheric ozone on the Δ_5-6_′ signal in tree-ring glucose

In Vienna, *[O_3_]* is measured at five sites. Complete timeseries for all sites are available since 1992. Intra-annually, the highest *[O_3_]* occurs during the period April to August (Fig. 6a) which can be expected to affect tree metabolism. Therefore, we calculated an April to August average timeseries for the Vienna region covering the period 1992 to 2020 (Fig. 6b, solid black line). We found that April to August *SD* and *rH* explain 59% of the timeseries variability (Fig. 6b, dashed black line, *p*<0.00001, *n*=29):

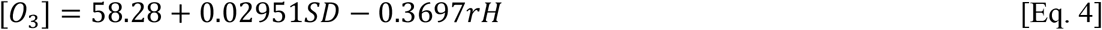

**Figure 6.**
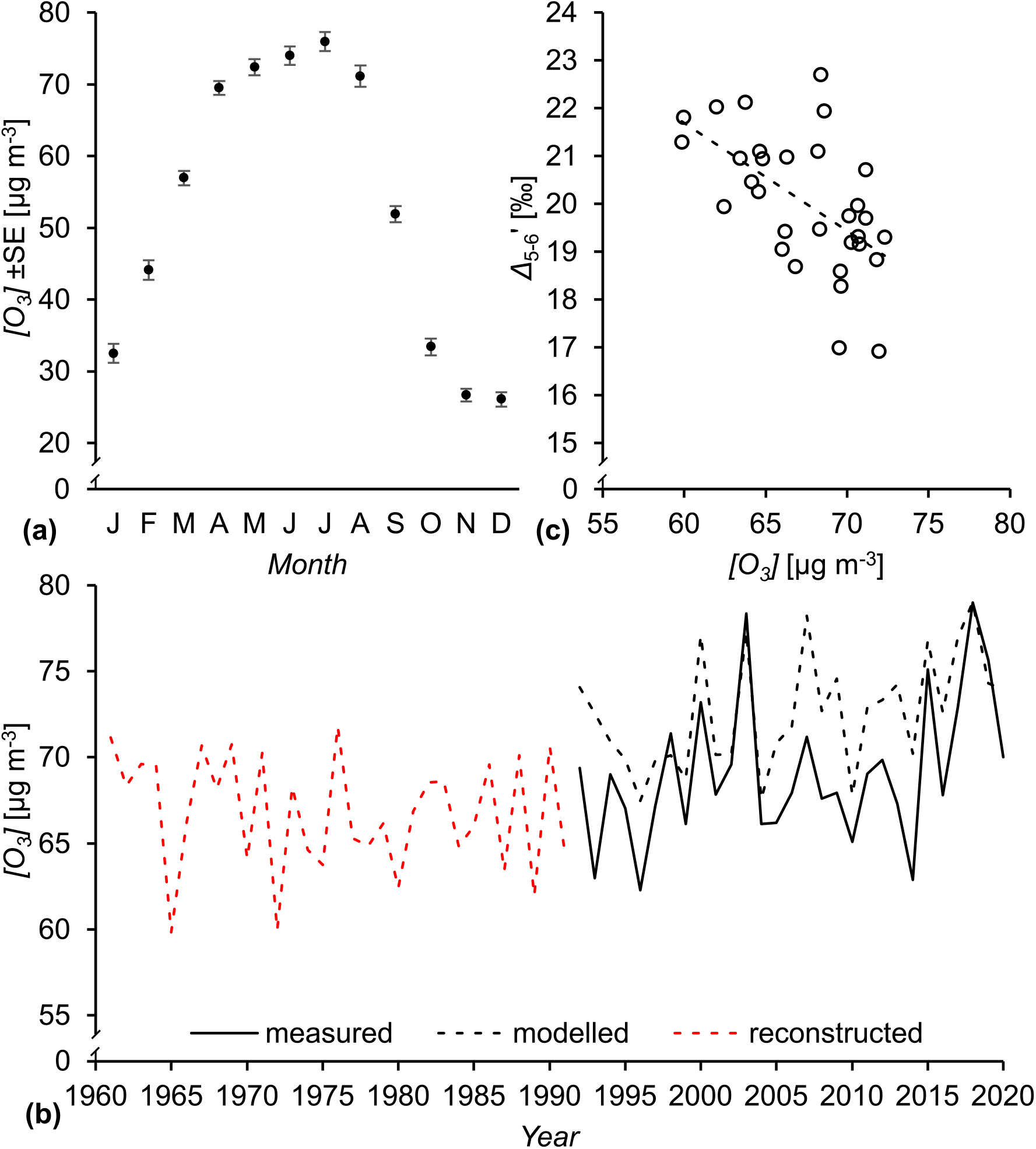
**(a)** Average monthly ozone concentrations, *[O_3_],* measured in Vienna over the period 1992 to 2020. **(b)** April to September *[O_3_]* in Vienna. Black solid line, measured *[O_3_];* black and red dashed lines, modelled and reconstructed *[O_3_]*, respectively, based on the relationship of *[O_3_]* with sunshine duration, *SD,* and relative humidity, *rH: [O_3_]*=58.28+0.02951*SD*-0.3697*rH*, *R^2^*=0.59, *p*<0.00001, *n*=29. **(c)** ^13^C discrimination at glucose C-5 and C-6, *Δ*_5-6_′, as function of *[O_3_]: Δ_5-6_*′=35.24-0.2258*[O_3_], R^2^*=0.33,*p*<0.001, *n*=31. Isotope data were measured on tree-ring glucose of *Pinus nigra* laid down from 1961 to 1995 at a dry site in the Vienna basin. *SD* and *rH* were measured at the climate station Hohe Warte (Vienna). *[O_3_]* was measured at five stations in Vienna (Stephansplatz, Laaer Berg, Hermannskogel, Hohe Warte, and Lobau).

Other studies report similar relationships (Felipe-Sotelo *et al*., 2006; Kovač-Andrić *et al*., 2009). Based on equation 4, we reconstructed *[O_3_]* for the Vienna region over the period 1961 to 1991 (Fig. 6b, dashed red line). We found that April to August *[O_3_]* (reconstructed for 1961 to 1991 and measured for 1992 to 1995) explains 33% of the *Δ*_5-6_′ timeseries variability (Fig. 6c, *p*<0.001, *n*=31):

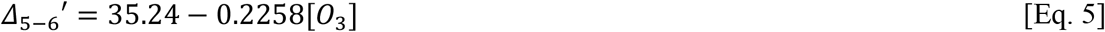

Accounting for measurement error, 75% of the variance in *Δ*_5-6_′ is explainable by modelling *(cf* Nilsson *et al*., 1996). Thus, *[O_3_]* explains 44% of the explainable variability in *Δ*_5-6_′ (33%/75%*100). This is substantial considering that measured and modelled *[O_3_]* timeseries merely exhibit 59% co-variability (Fig. 6b, solid and dashed black line). In summary, these results corroborate the proposed negative relationship between ozone stress and the *Δ*_5-6_′ signal (see ‘Ecophysiological effects’), and, by extension, the theoretical framework developed above.

### Implications of the theory

#### Signal propagation at the level of TPI in the cytosol of leaves

The post-Rubisco signal at glucose C-5 and C-6 is independent of a signal at glucose C-1 and C-2 (Fig. 4; Wieloch *et al*., 2018). That is, substantial signal propagation from C-5 and C-6 to C-2 and C-1 is not supported by the data. This is surprising for the following reason. Transmission of the *Δ*_5-6_′ signal from its origin, the lower end of the glycolytic pathway, into carbohydrate metabolism occurs via GAP (Fig. 1). Hence, signal independency requires negligible conversion of GAP (a precursor of glucose C-4 to C-6) to DHAP (a precursor of glucose C-1 to C-3) via leaf-cytosolic TPI (Figs. 1–2). Since TPI is often referred to as the prime example for the efficiency of enzyme catalysis, one would expect full equilibration of GAP and DHAP and inherent isotope signals. This view, however, is based on *in vitro* measurements of TPI kinetics. The following mechanisms may explain the apparent lack of equilibration and signal propagation *in vivo*.

In the light, chloroplast export of DHAP is favoured by the equilibrium position of stromal TPI, which is strongly on the side of DHAP (Walker, 1976; Knowles and Albery, 1977). Sharkey and Weise (2012) calculated there should be over 20 times more DHAP than GAP at equilibrium (Meyerhof and Junowicz-Kocholaty, 1943; Bassham and Krause, 1969). The substrate affinities of the triose phosphate translocator for DHAP and GAP are similar at *K_m_* = 0.13 mM and *K_m_*=0.08 mM, respectively (Fliege *et al*., 1978). Thus, DHAP and GAP will be transported according to their concentrations. That is, 20 times more DHAP will be exported from chloroplasts to the cytosol. However, synthesis of fructose 1,6-bisphosphate uses DHAP and GAP at a 1:1 ratio. This may keep the concentration of leaf-cytosolic GAP low. Flux of GAP into glycolysis and processes consuming glycolytic intermediates will additionally contribute to low cytosolic GAP concentrations. This may restrict the GAP to DHAP back conversions. Furthermore, numerous common metabolites inhibit TPI competitively (Anderson, 1971; Grüning *et al*., 2014; Flügel *et al*., 2017; Li *et al*., 2019). In addition, the activity of cytosolic TPI decreases significantly upon treatment with reactive oxygen species, especially H2O2 (Lopez-Castillo *et al*., 2016). Thus, during active photosynthesis, a lack of isomeric and isotopic equilibrium between leaf-cytosolic GAP and DHAP is conceivable. This would block the propagation of ^13^C signals in GAP to DHAP and enable independent ^13^C signals in *Δ*_5-6_′ and *Δ*_1-2_′ as observed.

#### Signal propagation to all glucose carbons via chloroplast metabolism

Cytosolic PEP, 2PGA, 3-phosphoglycerate, 1,3-bisphosphoglycerate, and GAP carry the *Δ*_5-6_′ signal from its putative origin (enolase, PEPC, PK, DAHPS) directly into C-5 and C-6 of hexose phosphates (Figs. 1–3). In addition, indirect signal propagation to C-1 to C-6 can be expected via import of 3-phosphoglycerate into chloroplasts (SI 1.2). A model of signal propagation described in SI 1 has several implications. First, observed *Δ*_5-6_′ signals at C-5 and C-6 are 6.625-fold larger than at C-1 to C-4 (SI 1.8). Second, the original signal as introduced at the level of enolase, PEPC, PK, DAHPS is 1.4-fold larger than the signal observed at glucose C-5 and C-6 (SI 1.9). Third, clustering of *Δ*_4_′ with the *Δ*_5_′ to *Δ*_6_ cluster (Fig. 4) can be explained by signal propagation via chloroplast metabolism (SI 1.10).

#### Signal propagation to other plant metabolites

Carbon flux changes around leaf-cytosolic enolase, PEPC, PK, and DAHPS proposedly introduce the *Δ*_5-6_′ signal. Hence, upstream derivatives of 2PGA carbons corresponding to glucose C-5 and C-6 will inherit the signal (Fig. 1). Compared to plant cellulose, the signal will be distinctly smaller in chloroplast derivatives, such as starch, and distinctly larger in leaf sucrose synthesised during the photoperiod (SI 1). This latter metabolite may be used to follow the *Δ*_5-6_′ signal on an hourly basis. Downstream derivatives of PEP carbons corresponding to glucose C-5 and C-6 will obtain an inverse *Δ*_5-6_′ signal. These differences may help to test the theory.

Turnover by PEPC, PK, and DAHPS can be expected to result in ^13^C-enriched PEP (see ‘Leaf-level enolase, PEPC, PK, and/or DAHPS introduce the *Δ*_5-6_′ signal’). Thus, our theory can help to explain the ^13^C/^12^C separation observed among plant compounds, specifically between ^13^C-enriched carbohydrates and ^13^C-depleted metabolites downstream of PK, PEPC, and DAHPS (Sharkey *et al*., 1991; Bathellier *et al*., 2017).

#### Implications for whole-molecule ^13^C/^12^C analysis

The *Δ*_5-6_′ signal has two components of variance (see ‘Ecophysiological effects’). One is inversely correlated with diffusion-Rubisco fractionation, the other is independent. Both components have implications for studies of plant carbon uptake and associated properties by whole-molecule ^13^C/^12^C analysis. The inversely correlated component removes the diffusion-Rubisco signal from glucose C-5 and C-6. In addition, this signal is absent at glucose C-4 (Wieloch *et al*., 2018). That is, three out of six glucose carbon positions lack the diffusion-Rubisco signal. Thus, whole-molecule ^13^C/^12^C analysis captures an attenuated diffusion-Rubisco signal and underestimates the variability of the original signal and associated physiological properties, such as *C*_i_*/C*_a_ and photosynthetic water-use efficiency.

The independent component of the *Δ*_5-6_′ signal weakens signal extractions from whole-molecule ^13^C/^12^C measurements because it constitutes pseudorandom noise with respect to diffusion-Rubisco fractionation. This may explain why models of whole-molecule diffusion-Rubisco fractionation as functions of environmental properties often suffer from low explanatory powers, *R^2^*≤0.5 (Barbour and Song, 2014). By contrast, intramolecular ^13^C/^12^C analysis resolves information about distinct ecophysiological processes; a fundamental conceptual advancement enabling more adequate modelling of the variability of plant carbon uptake and associated environmental/developmental controls.

### Tracking carbon allocation in other biological organisms

Whole-molecule ^13^C/^12^C analysis enables assessments of plant carbon uptake (Farquhar *et al*., 1982). According to theory reported here, intramolecular ^13^C/^12^C analysis enables additional assessment of downstream carbon allocation in actively photosynthesising leaves. This includes carbon flux into the anaplerotic, shikimate, MEP, and fatty acid biosynthesis pathways, and mitochondrial respiration (Fig. 1). Intramolecular ^13^C signals are governed by a small set of physicochemical principles that apply generally (Schmidt *et al*., 2015). Thus, intramolecular ^13^C/^12^C analysis can be expected to enable retrospective assessment of carbon allocation in any biological organism including, for instance, disease-related shifts.

### Utility of the *Δ*_5-6_′ signal

Laboratory experiments are limited to short timescales and in their capabilities to reproduce complex natural systems. Manipulation experiments on natural systems are limited to timescales of years and may suffer from spurious effects due to unnatural step changes in ambient conditions. By contrast, tree-ring analysis offers extensive temporal, spatial, species, and genotype coverage of natural systems that have not been subjected to unnatural step changes.

Proposedly, the *Δ*_5-6_′ signal reports flux into the anaplerotic pathway including CO_2_ uptake by PEPC (Figs. 1, 3, and 5–6). In addition, it may report flux into mitochondrial respiration (SI 3). Thus, signal analysis may enable a better understanding of plant and ecosystem carbon balances including the so-called CO_2_ fertilisation effect.

Furthermore, intramolecular ^13^C/^12^C analysis not only enables analysis of carbon uptake-environment relationships but also of carbon allocation-environment relationships (Figs. 5–6) and thus more comprehensive assessments of flux-level plant performance. For instance, atmospheric CO_2_ and ozone concentrations have increased over recent years (Fig. 6b). Under business-as-usual scenarios, this will continue over the next decades (Turnock *et al*., 2018). While CO_2_ promotes leaf photosynthesis and net primary productivity, ozone has the very opposite effect (Ainsworth *et al*., 2012; IPCC, 2014). When this highly reactive chemical enters plant leaves through stomata, it causes harm to structure and function and leads to major rearrangements in carbon metabolism. While ozone decreases carbon fixation, it increases carbon allocation to costly maintenance and repair processes (Dizengremel, 2001; Ainsworth *et al*., 2012). This includes increased carbon flux into the anaplerotic and shikimate pathway, and this resource investment is likely recorded in the *Δ*_5-6_′ signal. Ozone tolerance varies among species with metabolic changes depending on the duration of ozone exposure (Fontaine *et al*., 2003; Ainsworth *et al*., 2012). Thus, the *Δ*_5-6_′ signal may support flux-level screenings for species/genotypes with the capacity to optimally adjust to prolonged ozone exposure (requires further investigation).

While glucose positions C-1 to C-3 preserve the VPD-dependent carbon uptake signal (Wieloch *et al*., 2018), this signal was removed from glucose C-5 and C-6 and replaced by an independent ozone-sensitive carbon allocation signal (Figs. 5–6). Thus, intramolecular ^13^C/^12^C analysis yields information about several environmental variables and may enable more powerful paleoenvironment reconstructions than whole-molecule analysis.

Lastly, sampling glucose at different developmental stages may enable the detection of shifts in carbon uptake and allocation over the lifespan of plants to better understand basic physiological processes such as plant senescence. In conclusion, intramolecular ^13^C/^12^C analysis opens up numerous new avenues of research within the plant and Earth sciences.

## Supplementary data

SI 1 Signal propagation to all glucose carbons via chloroplast metabolism and implications

SI 1.1 Signal propagation upon reimport of cytosolic metabolites into chloroplasts

SI 1.2 Predominant import molecules

SI 1.3 Predominant export molecules

SI 1.4 Signal import into chloroplasts

SI 1.5 Signal dilution and partial signal loss

SI 1.6 Stromal signal redistribution

SI 1.7 Signal size in stromal triose phosphates

SI 1.8 Signal size in cytosolic hexose phosphates

SI 1.9 Original size of the *Δ*_5-6_′ signal

SI 1.10 Signal propagation can explain the clustering of the *Δ*_4_ to *Δ6* timeseries

SI 1.11 The role of chloroplast starch in signal propagation

SI 1.12 Signal import into chloroplasts via cytosolic 1,3BPGA, 2PGA, and PEP

SI 1.13 Assumptions

SI 2 Correction for ^13^C signal redistribution by triose phosphate cycling in tree-ring cells

SI 3 Alternating substrate supply to oxidative phosphorylation may contribute to the Δ_5-6_’ signal

SI 4 Carbon flux into fatty acid biosynthesis in illuminated photosynthetic tissue

SI 5 Potential influence of environmental and developmental variables on the Δ_5-6_’ signal

SI 6 Intramolecular isotope effects in response to ozone

## Acknowledgments

We thank Lisa Wingate (INRA, Ephyse Research Unit, Bordeaux, France), Richard Leegood (University of Sheffield, UK), and Nicole Linka (Heinrich Heine University Düsseldorf, Germany) for helpful comments. T.W.’s and J.S.’s research was funded by the Swedish Research Council VR (2013-05219, 2018-04456), the Knut and Alice Wallenberg Foundation (“NMR for Life” facility and grant 2015.0047), and the Kempe foundations. T.D.S.’s research was funded by U.S. Department of Energy (DE-FG02-91ER2002). Partial salary support for T.D.S. came from Michigan AgBioResearch.

## Author Contribution

T.W. conceived the study with input from T.D.S., J.S., and R.A.W. T.W. led the research. T.W. devised the physiological part of the study with input from T.D.S., R.A.W., and J.S. T.W. devised the isotope part of the study with input from R.A.W., T.D.S., and J.S. T.W. analysed the data with input from R.A.W. T.W. wrote the manuscript with input from T.D.S., J.S., and R.A.W.

## Data Availability

The data that support the findings of this study have been published previously by Wieloch *et al.* (2018) and Saurer *et al.* (1995).

