## Supplementary information for "Intramolecular carbon isotope signals reflect metabolite allocation in plants"

***SI 1 Signal propagation to all glucose carbons via chloroplast metabolism and implications***

<u>SI 1.1 Signal propagation upon reimport of cytosolic metabolites into chloroplasts.</u> According to our theory, cytosolic PEP, 2PGA, 3PGA, 1,3BPGA, and GAP carry the $\Delta_{5\text{-}6}'$ signal from its putative origin (enolase, PEPC, PK, DAHPS) directly into hexose phosphate C-5 and C-6 (Fig. 1). If these metabolites were reimported into chloroplasts at high rates, significant propagation of the $\Delta_{5\text{-}6}'$ signal to other carbon positions may occur. However, $\Delta_5'$ and $\Delta_6'$ are not related with $\Delta_1'$ to $\Delta_3'$, and only weakly related with $\Delta_4'$ (Fig. 4). Thus, substantial $\Delta_{5\text{-}6}'$-signal propagation to other carbon positions is not supported by the data. That is, our theory requires negligible signal propagation via chloroplast metabolism. This requirement is met by signal dilution and partial signal loss as follows.

<u>SI 1.2 Predominant import molecules.</u> Transport of 3PGA, GAP, DHAP, and $P_i$ across the inner envelop of chloroplasts is mediated by the triose phosphate translocator (Heber and Heldt, 1981; Linka and Weber, 2010), with a strong preference for a 1:1 counter-exchange of substrates (unidirectional transport rates are 2 to 3 orders of magnitude lower; Fliege *et al.*, 1978; Mourioux and Douce, 1981; Neuhaus and Maaß, 1996). That is, for 1 molecule exported from chloroplasts, 1 cytosolic molecule is imported. Since the triose phosphate translocator exhibits no pronounced preference for 3PGA, GAP, DHAP, or $P_i$ (Fliege *et al.*, 1978; Kammerer *et al.*, 1998), import largely depends on relative cytosolic substrate concentrations. Data on substrate concentrations in the cytosol of illuminated leaves are scarce. In *Spinacia oleracea*, 3PGA levels of ≈120 nmol mg $Chl^{-1}$ and DHAP levels of ≈10 nmol mg $Chl^{-1}$ were reported (photoperiod averages; Gerhardt *et al.*, 1987). In *Phaseolus vulgaris*, $P_i$ levels of ≈350 nmol mg $Chl^{-1}$ were reported (at light-saturating conditions; Sharkey and Vanderveer, 1989). Hence, levels of DHAP are an order of magnitude lower than levels of 3PGA, and $P_i$. Cytosolic levels of GAP can be expected to be at least an order of magnitude smaller than levels of DHAP (see main text). Thus, (i) cytosolic 3PGA, and $P_i$ are the predominant import molecules, and (ii) substantial signal transmission into chloroplasts by cytosolic GAP can be excluded (cytosolic DHAP, and GAP are not considered further).

SI 1.3 Predominant export molecules. In the light, stromal $P_i$ levels are low (Sharkey and Vanderveer, 1989), and export of 3PGA is believed to be restricted due to the chloroplast-to-cytosol pH gradient (Lilley *et al.*, 1977; Heldt *et al.*, 1978; Fliege *et al.*, 1978; Flügge *et al.*, 1983). Thus, triose phosphates are the predominant export molecules (stromal $P_i$, and 3PGA are not considered further).

SI 1.4 Signal import into chloroplasts. Reported levels of cytosolic $P_i$ exceed 3PGA levels by about a factor of 3 (see above). Therefore, gross export of 4 stromal triose phosphates involves reimport of 3 molecules $P_i$, and 1 molecule 3PGA. This counter-exchange results in net export of 3 molecules triose phosphate. The reimported molecule 3PGA carries the $\Delta_{5\text{-}6}'$ signal into chloroplasts. Upon 3PGA reduction, the signal enters the stromal triose phosphate pool.

SI 1.5 Signal dilution and partial signal loss. Net export of 3 molecules triose phosphate requires net fixation of 9 carbon atoms by Rubisco. In $C_3$ plants, ratios of Rubisco oxygenation to carboxylation of $v_O/v_C$=0.4 are common (Sharkey, 1988). Per two oxygenations, 1 molecule $CO_2$ is released. Thus, net fixation of 9 carbon atoms requires an actual fixation of 11.25 molecules $CO_2$ (I. net fixation=$v_C$-$v_O$/2, II. $v_O$=0.4$v_C$, II. in I. $v_C$=net fixation/0.8) and entails photorespiratory loss of 2.25 molecules $CO_2$. Newly fixed carbon is free of the $\Delta_{5\text{-}6}'$ signal and leads to signal dilution in the stromal triose phosphate pool. In addition, photorespiratory $CO_2$ release results in partial signal loss from metabolism (see assumptions below).

SI 1.6 Stromal signal redistribution. *De novo* synthesis of three molecules triose phosphate involves 11.25 carboxylations and 4.5 oxygenations of ribulose 1,5-bisphosphate (see above). Thus, 15.75 molecules ribulose 1,5-bisphosphate need to be regenerated, and this involves redistributing carbon of 26.25 molecules triose phosphate (15.75*5/3). In addition, photorespiration involves carbon redistribution. Carbon redistribution can be expected to cause $\Delta_{5\text{-}6}'$-signal redistribution over all carbon positions of stromal triose phosphate. This is supported by the observation of an even intramolecular $^{14}C$ distribution in starch glucose after 45 minutes labelling time (Gibbs and Kandler, 1957).

SI 1.7 Signal size in stromal triose phosphates. In the steady state, the $\Delta_{5\text{-}6}'$ signal enters cytosolic 3PGA by two pathways. First, the signal is transmitted directly from its source (enolase, PEPC, PK, and/or DHAPS; transmission from the bottom). Second, the signal is transmitted via triose phosphates exported from chloroplasts (due to 3PGA reimport into chloroplasts; transmission from the top). Thus, all carbon positions of cytosolic 3PGA can carry fractions of the $\Delta_{5\text{-}6}'$ signal. We assume that imported 3PGA carries a signal of size 1 into the stromal triose phosphate pool, thereby adding 3 carbon atoms to the pool. *De novo* synthesis of 3 molecules triose phosphate adds another 11.25 carbon atoms to the pool. After carbon redistribution (see above), 14.25 carbon atoms each contain a signal of size 1/14.25. Due to photorespiratory loss of 2.25 carbons, a fraction of 2.25/14.25 of the signal is lost from metabolism.

SI 1.8 Signal size in cytosolic hexose phosphates. Exported triose phosphates contain a signal of size 1/14.25 per carbon position. All carbon positions of cytosolic hexose phosphate inherit this signal. Hexose phosphate carbons 4 to 6 exhibit a signal of size 1 because cytosolic 3PGA and GAP are in equilibrium (see main text). Hexose phosphate C-5 and C-6 contain a signal of size (1-1/14.25)/2 each. Thus, signals at C-5 and C-6 are 6.625-fold larger than signals at C-1 to C-4 ((1-1/14.25)/2)/(1/14.25).

SI 1.9 Original size of the $\Delta_{5\text{-}6}'$ signal. The largest fraction of the original $\Delta_{5\text{-}6}'$ signal, as introduced at the level of enolase, PEPC, PK, and DAHPS, is transmitted to C-5 and C-6 of cytosolic hexose phosphate. However, smaller fractions enter C-1 to C-4 of cytosolic hexose phosphate and photorespiratory $CO_2$ via chloroplast metabolism. Thus, the size of the original signal exceeds the size of the apparent signal at tree-ring glucose C-5 and C-6, and we will now estimate the size of the original signal. Cytosolic hexose phosphate C-5 and C-6 together carry a signal of size 1-1/14.25. Cytosolic hexose phosphate C-1 to C-4 carry a signal of size 1/14.25 each. Photorespiration causes signal loss per hexose phosphate of size 2.25/14.25*2/3. Thus, the original signal has the size 1.3 ($s_O$=(1-1/14.25)+4/14.25+2.25/14.25*2/3), i.e., it is 1.4-fold larger than the apparent signal at hexose phosphate C-5 and C-6 ($s_O$/(1-1/14.25)).

SI 1.10 Signal propagation can explain the clustering of the $\Delta_4'$ to $\Delta_6'$ timeseries. In figure 4, $\Delta_1'$ to $\Delta_3'$ form a cluster. Wieloch *et al.* (2018) argue that variability in this cluster is explained by diffusion-Rubisco fractionation. Moreover, the separation between $\Delta_3'$ and $\Delta_1'$ and $\Delta_2'$ is explained by an additional post-Rubisco signal in $\Delta_1'$ and $\Delta_2'$ which is partly correlated with the diffusion-Rubisco signal (Wieloch *et al.*, 2018). The $\Delta_1'$ to $\Delta_3'$ cluster is independent of the $\Delta_4'$ to $\Delta_6'$ cluster (Fig. 4). Wieloch *et al.* (2018) showed evidence for the absence of the diffusion-Rubisco signal at C-4. In the main text, we argue the diffusion-Rubisco signal is absent at C-5 and C-6. Thus, propagation of the $\Delta_{5\text{-}6}$' signal to hexose phosphate C-4 via chloroplast metabolism can explain $\Delta_4$' to $\Delta_6$' cluster formation.

SI 1.11 The role of chloroplast starch in signal propagation. Part of the $\Delta_{5\text{-}6}$' signal is transmitted to the stromal triose phosphate pool via reimport of cytosolic 3PGA into chloroplasts. Stromal triose phosphates are the precursors of both chloroplast starch and cytosolic sucrose synthesised during the photoperiod. Thus, the two carbon pools both obtain parts of the $\Delta_{5\text{-}6}$'-signal. However, this signal partitioning is reversed during synthesis of tree-ring glucose and plant cellulose which utilises both carbon sources (see assumptions below). Thus, separate consideration of signal propagation via starch is not required here.

SI 1.12 Signal import into chloroplasts via cytosolic 1,3BPGA, 2PGA, and PEP. Transport of 1,3BPGA across the inner envelop of chloroplasts is generally not discussed. There are two reasons why it is unlikely to occur at significant rates. First, *in vivo* 1,3BPGA concentrations can be expected to be low due to the equilibrium positions of reactions catalysed by p-GAPDH and PGK: $K_{\text{p-GAPDH}}$=0.63, $K_{\text{PGK}}$=3180 (Figs. 1, and 2; Meyerhof and Oesper, 1947; Bücher, 1947; Bassham and Krause, 1969). Second, the triose phosphate translocator efficiently transports doubly charged anions in the form of three-carbon molecules with one terminal phosphate group (Heldt *et al.*, 1978; Flügge *et al.*, 1983). By contrast, 1,3BPGA has 4 negative charges and two terminal phosphate groups which should impede its transport. Similarly, there are two reasons why 2PGA transport is unlikely to occur at significant rates. First, *in vivo* 2PGA concentrations can be expected to be low due to the equilibrium positions of the reactions catalysed by PGM and enolase: $K_{\text{PGM}}$=0.095, $K_{\text{enolase}}$=4 (Figs. 1 and, 3; Utter and Werkman, 1942; Wold and Ballou, 1957; Bassham and Krause, 1969). Second, the affinity of the triose phosphate translocator for 2PGA is at least an order of magnitude lower than for DHAP, GAP, and $P_i$ (Fliege *et al.*, 1978). Thus, transmission of the $\Delta_{5\text{-}6}$' signal into chloroplast metabolism via 1,3BPGA and 2PGA can be expected to be negligible. Reimport of $^{13}$C signals in PEP into chloroplasts occurs by the PEP/$P_i$ translocator. However, PEP is biochemically separated from the stromal triose phosphate pool due to the lack of enolase in leaf chloroplasts (see main text). Thus, direct signal propagation from PEP to stromal triose phosphates and derivatives is not feasible.

SI 1.13 Assumptions. Our discussion about signal propagation makes several implicit assumptions. First, photorespiration is a closed cycle recovering as much carbon as 3PGA as possible. Abstraction of photorespiratory metabolites for other processes would cause signal loss from carbohydrate metabolism. Second, re-fixation of photorespiratory $CO_2$ is not considered but would compensate photorespiratory signal loss. Third, utilisation of metabolites involved in $\Delta_{5\text{-}6}'$-signal signal transmission to tree-ring glucose by processes other than tree-ring glucose synthesis is not considered but would entail further signal loss from carbohydrate metabolism. Refined modelling of signal propagation should consider such processes carefully. Nevertheless, we believe our model is a good first approximation of signal propagation and its implications.

### *SI 2 Correction for $^{13}$C signal redistribution by triose phosphate cycling in tree-ring cells*

Sucrose translocated from leaf to tree-ring cells is converted into different hexose derivatives, the precursors of tree-ring glucose. 40 to 50% of these derivatives participate in futile carbon cycling with triose phosphates (Fig. 1). In tree-ring cells, the triose phosphates glyceraldehyde 3-phosphate (glucose C-4 to C-6) and dihydroxyacetone phosphate (glucose C-1 to C-3) are in complete or close to complete equilibrium (see 'Exclusion of the tree-ring cell as origin of the $\Delta_{5\text{-}6}'$ signal' in the main text). As a result, 20 to 25% of any $^{13}$C signals present at a specific carbon position in the total hexose derivative pool arrive at the symmetry-related carbon position in tree-ring glucose (C-1 and C-6, C-2 and C-5, and C-3 and C-4). Wieloch *et al.* (2018) reported a model of $^{13}$C signal redistribution by triose-phosphate cycling (TPC) which can be used to remove TPC-related effect. This restores the distribution of $^{13}$C signals as introduced at the leaf-level.

### *SI 3 Alternating substrate supply to oxidative phosphorylation may contribute to the $\Delta_{5\text{-}6}'$ signal*

In illuminated photosynthetic tissue, photorespiration and mitochondrial respiration are believed to act in concert, jointly ensuring optimal cell functioning under varying environmental conditions (Hurry *et al.*, 2005). Alternating substrate supply to oxidative phosphorylation may contribute to the component of the $\Delta_{5\text{-}6}'$ signal that is inversely correlated with diffusion-Rubisco fractionation. Normally, mitochondrial oxidation of photorespiratory glycine is believed to support oxidative phosphorylation (Krömer, 1995; Gardeström and Igamberdiev, 2016). At low photorespiratory rates however, pyruvate can be expected to fill in for glycine as respiratory substrate as follows (Gemel and Randall, 1992).

Pyruvate enters the tricarboxylic acid cycle via the mitochondrial pyruvate dehydrogenase complex (PDC). In the light, mitochondrial PDC is partly inhibited due to reversible phosphorylation (Budde and Randall, 1990; Gemel and Randall, 1992; Tovar-Mendez *et al.*, 2003; Tcherkez *et al.*, 2005). Phosphorylation is mediated by a specific kinase which is activated by a high ATP/ADP ratio and ammonium (Tovar-Mendez *et al.*, 2003). In addition, NADH at physiological concentrations inhibits PDC allosterically (Tovar-Mendez *et al.*, 2003). Photorespiration produces ammonium and NADH inside mitochondria. A significant fraction of this NADH is believed to feed into oxidative phosphorylation supplying the mitochondrial matrix with ATP (Krömer, 1995; Gardeström and Igamberdiev, 2016). Accordingly, leaf protoplasts exhibit substantially increased mitochondrial $NADH/NAD^+$ and ATP/ADP ratios under photorespiration (Gardeström and Wigge, 1988; Wigge *et al.*, 1993; Igamberdiev *et al.*, 2001; Igamberdiev and Gardeström, 2003; Gardeström and Igamberdiev, 2016). Thus, reduced photorespiration can be expected to result in PDC activation (Gardeström and Igamberdiev, 2016), and this is supported by the following two observations. First, *in vivo* PDC activity increased in illuminated *Pisum sativum* seedlings at high ambient $CO_2$ and even further at additionally low ambient $O_2$ (Budde and Randall, 1990). Second, photorespiratory inhibitors increase PDC activity in detached illuminated leaves of *Pisum sativum* (Gemel and Randall, 1992). PDC catalyses the conversion of pyruvate to acetyl-CoA. *In vivo*, PK can be expected to govern pyruvate supply to PDC because PEP to pyruvate flux via PK strongly exceeds malate to pyruvate flux via malic enzyme (Plaxton and Podestá, 2006). Thus, a negative correlation between photorespiration and carbon flux through PK can be expected, and this may remove the diffusion-Rubisco signal from glucose C-5 and C-6 as follows.

For all practical purposes, $O_2$ levels in chloroplasts of terrestrial $C_3$ plants can be considered constant (Ligeza *et al.*, 1997). Thus, the chloroplast $CO_2/O_2$ ratio is governed by the $CO_2$ component. At high levels of chloroplast $CO_2$, diffusion-Rubisco fractionation results in the synthesis of relatively $^{13}C$-depleted glucose (Farquhar *et al.*, 1982; Evans *et al.*, 1986). Concomitantly high $CO_2/O_2$ inhibits photorespiration and stimulates ATP-consuming sucrose synthesis (Gardeström and Igamberdiev, 2016). This can be expected to result in high flux through PK to meet high cytosolic ATP demands (Gardeström and Igamberdiev, 2016). Carbon turnover by PK can be expected to cause $^{13}C$ enrichment in the remaining substrate PEP at carbon positions corresponding to glucose C-5 and C-6 (see main text). Thus, increased $^{13}C$ enrichment at high flux rates can be expected to counteract $^{13}C$-depletions by diffusion-Rubisco fractionation.

***SI 4 Carbon flux into fatty acid biosynthesis in illuminated photosynthetic tissue***

In illuminated photosynthetic tissue of *Arabidopsis thaliana*, fatty acid biosynthesis occurs at a rate of 2.3 µmol C mg $Chl^{-1}$ $h^{-1}$ (Bao *et al.*, 2000). In wild-type plants, chlorophyll contents of 0.25 g Chl $m^{-2}$ are common (Li *et al.*, 2019). Using this value, 2.3 µmol C mg $Chl^{-1}$ $h^{-1}$ can be converted to 0.16 µmol C $m^{-2}$ $s^{-1}$. In leaves of *Arabidopsis thaliana*, net $CO_2$ assimilation rates of 7.5 µmol C $m^{-2}$ $s^{-1}$ are common at ambient $CO_2$ concentrations of 380 ppm and light-saturating conditions (Tanaka *et al.*, 2013). Thus, carbon flux into fatty acid biosynthesis reported by Bao *et al.* (2000) is estimated to account for ≈2% of net $CO_2$ assimilation.

***SI 5 Potential influence of environmental and developmental variables on the $\Delta_{5\text{-}6}'$ signal***

In general, the $\Delta_{5\text{-}6}'$ signal might be under developmental control. However, formation of *Pinus nigra* tree rings analysed by Wieloch *et al.* (2018) occurred when the trees had reached the canopy. On average, tree-ring width measurements of these samples reach back to 1865 (range: 1840 to 1918), and $^{13}C/^{12}C$ measurements start at 1961. Thus, in the present case, the $\Delta_{5\text{-}6}'$ signal can be expected to be under environmental rather than developmental control.

***SI 6 Intramolecular isotope effects in response to ozone***

Let $\Delta\Delta$ and $\Delta\Delta_i$ denote whole-molecule and intramolecular shifts of $^{13}C$ discrimination, respectively, with *i* denoting intramolecular carbon positions in glucose. $\Delta\Delta$ and $\Delta\Delta_i$ are related as $\Delta\Delta=\Sigma\Delta\Delta_i/n$ where *n* denotes the number of intramolecular carbon positions. According to the discussion about signal propagation, $\Delta\Delta_5$ and $\Delta\Delta_6$, denoted *x*, are 6.625-fold larger than $\Delta\Delta_1$ to $\Delta\Delta_4$, denoted *y*; $x=6.625y$. On average, post-Rubisco fractionation caused $\Delta\Delta$ decreases of ≈-1.98 ± 0.58SE ‰ in leaf and stem cellulose of *Betula pendula* in response to ozone (Fig. 5). Solving $\Delta\Delta=(4y+2x)/6\approx-1.98 \pm 0.58$SE ‰ for *x* and *y* gives $x=\Delta\Delta_5=\Delta\Delta_6\approx-4.56 \pm 1.34$SE ‰ and $y=\Delta\Delta_1=\Delta\Delta_2=\Delta\Delta_3=\Delta\Delta_4\approx-0.69 \pm 0.20$SE ‰, respectively.

*References*

**Bao X, Focke M, Pollard M, Ohlrogge J**. 2000. Understanding *in vivo* carbon precursor supply for fatty acid synthesis in leaf tissue. The Plant Journal **22**, 39–50.

**Bassham JA, Krause GH**. 1969. Free energy changes and metabolic regulation in steady-state photosynthetic carbon reduction. Biochimica et Biophysica Acta **189**, 207–221.

**Bücher T**. 1947. Über ein phosphatübertragendes Gärungsferment. Biochimica et Biophysica Acta **1**, 292–314.

**Budde RJ, Randall DD**. 1990. Pea leaf mitochondrial pyruvate dehydrogenase complex is inactivated *in vivo* in a light-dependent manner. Proceedings of the National Academy of Sciences of the United States of America **87**, 673–676.

**Evans JR, Farquhar GD, Sharkey TD, Berry JA**. 1986. Carbon isotope discrimination measured concurrently with gas exchange to investigate $CO_2$ diffusion in leaves of higher plants. Australian Journal of Plant Physiology **13**, 281–292.

**Farquhar GD, O'Leary MH, Berry JA**. 1982. On the relationship between carbon isotope discrimination and the intercellular carbon dioxide concentration in leaves. Australian Journal of Plant Physiology **9**, 121–137.

**Fliege R, Flügge UI, Werdan K, Heldt HW**. 1978. Specific transport of inorganic phosphate, 3-phosphoglycerate and triosephosphates across the inner membrane of the envelope in spinach chloroplasts. Biochimica et Biophysica Acta **502**, 232–247.

**Flügge UI, Gerber J, Heldt HW**. 1983. Regulation of the reconstituted chloroplast phosphate translocator by an $H^+$ gradient. Biochimica et Biophysica Acta **725**, 229–237.

**Gardeström P, Igamberdiev AU**. 2016. The origin of cytosolic ATP in photosynthetic cells. Physiologia Plantarum **157**, 367–379.

**Gardeström P, Wigge B**. 1988. Influence of photorespiration on ATP/ADP ratios in the chloroplasts, mitochondria, and cytosol, studied by rapid fractionation of barley (*Hordeum vulgare*) protoplasts. Plant Physiology **88**, 69–76.

**Gemel J, Randall DD**. 1992. Light regulation of leaf mitochondrial pyruvate dehydrogenase complex: Role of photorespiratory carbon metabolism. Plant Physiology **100**, 908–914.

**Gerhardt R, Stitt M, Heldt HW**. 1987. Subcellular metabolite levels in spinach leaves: Regulation of sucrose synthesis during diurnal alterations in photosynthetic partitioning. Plant Physiology **83**, 399–407.

**Gibbs M, Kandler O**. 1957. Asymmetric distribution of $C^{14}$ in sugars formed during photosynthesis. Proceedings of the National Academy of Sciences of the United States of America **43**, 446–451.

**Heber U, Heldt HW**. 1981. The chloroplast envelope: structure, function, and role in leaf metabolism. Annual Review of Plant Physiology **32**, 139–168.

**Heldt HW, Flügge UI, Fliege R**. 1978. The influence of illumination on the transport of 3-phosphoglycerate across the chloroplast envelope. In: Azzone GF, ed. The Proton and Calcium Pumps. New York: Elsevier/North-Holland Biomedical Press, 105–114.

**Hurry V, lgamberdiev AU, Keerberg O, Pärnik T, Atkin OK, Zaragoza-Castells J, Gardeström P**. 2005. Respiration in photosynthetic cells: Gas exchange components, interactions with photorespiration and the operation of mitochondria in the light. In: Lambers H, Ribas-Carbo M, eds. Advances in Photosynthesis and Respiration. Plant Respiration: From Cell to Ecosystem. Dordrecht: Springer, 43–61.

**Igamberdiev AU, Bykova NV, Lea PJ, Gardeström P**. 2001. The role of photorespiration in redox and energy balance of photosynthetic plant cells: A study with a barley mutant deficient in glycine decarboxylase. Physiologia Plantarum **111**, 427–438.

**Igamberdiev AU, Gardeström P**. 2003. Regulation of NAD- and NADP-dependent isocitrate dehydrogenases by reduction levels of pyridine nucleotides in mitochondria and cytosol of pea leaves. Biochimica et Biophysica Acta **1606**, 117–125.

**Kammerer B, Fischer K, Hilpert B, Schubert S, Gutensohn M, Weber A, Flügge UI**. 1998. Molecular characterization of a carbon transporter in plastids from heterotrophic tissues: the glucose 6-phosphate/phosphate antiporter. Plant Cell **10**, 105–117.

**Krömer S**. 1995. Respiration during photosynthesis. Annual Review of Plant Physiology and Plant Molecular Biology **46**, 45–70.

**Li J, Weraduwage SM, Preiser AL, Tietz S, Weise SE, Strand DD, Froehlich JE, Kramer DM, Hu J, Sharkey TD**. 2019. A cytosolic bypass and G6P shunt in plants lacking peroxisomal hydroxypyruvate reductase. Plant Physiology **180**, 783–792.

**Ligeza A, Tikhonov AN, Subczynski WK**. 1997. *In situ* measurements of oxygen production and consumption using paramagnetic fusinite particles injected into a bean leaf. Biochimica et Biophysica Acta **1319**, 133–137.

**Lilley RM, Chon CJ, Mosbach A, Heldt HW**. 1977. The distribution of metabolites between spinach chloroplasts and medium during photosynthesis *in vitro*. Biochimica et Biophysica Acta **460**, 259–272.

**Linka N, Weber APM**. 2010. Intracellular metabolite transporters in plants. Molecular Plant **3**, 21–53.

**Meyerhof O, Oesper P**. 1947. The mechanism of the oxidative reaction in fermentation. Journal of Biological Chemistry **170**, 1–22.

**Mourioux G, Douce R**. 1981. Slow passive diffusion of orthophosphate between intact isolated chloroplasts and suspending medium. Plant Physiology **67**, 470–473.

**Neuhaus H, Maaß U**. 1996. Unidirectional transport of orthophosphate across the envelope of isolated cauliflower-bud amyloplasts. Planta **198**, 542–548.

**Plaxton WC, Podestá FE**. 2006. The functional organization and control of plant respiration. Critical Reviews in Plant Sciences **25**, 159–198.

**Sharkey TD**. 1988. Estimating the rate of photorespiration in leaves. Physiologia Plantarum **73**, 147–152.

**Sharkey TD, Vanderveer PJ**. 1989. Stromal phosphate concentration is low during feedback limited photosynthesis. Plant Physiology **91**, 679–684.

**Tanaka Y, Sugano SS, Shimada T, Hara-Nishimura I**. 2013. Enhancement of leaf photosynthetic capacity through increased stomatal density in *Arabidopsis*. New Phytologist **198**, 757–764.

**Tcherkez G, Cornic G, Bligny R, Gout E, Ghashghaie J**. 2005. *In vivo* respiratory metabolism of illuminated leaves. Plant Physiology **138**, 1596–1606.

**Tovar-Mendez A, Miernyk JA, Randall DD**. 2003. Regulation of pyruvate dehydrogenase complex activity in plant cells. European Journal of Biochemistry **270**, 1043–1049.

**Utter MF, Werkman CH**. 1942. Dissimilation of phosphoglyceric acid by *Escherichia coli*. Biochemical Journal **36**, 485–493.

**Wieloch T, Ehlers I, Yu J, Frank D, Grabner M, Gessler A, Schleucher J**. 2018. Intramolecular $^{13}C$ analysis of tree rings provides multiple plant ecophysiology signals covering decades. Scientific Reports **8**, 5048.

**Wigge B, Krömer S, Gardeström P**. 1993. The redox levels and subcellular distribution of pyridine nucleotides in illuminated barley leaf protoplasts studied by rapid fractionation. Physiologia Plantarum **88**, 10–18.

**Wold F, Ballou CE**. 1957. Studies on the enzyme enolase. I. Equilibrium studies. Journal of Biological Chemistry **227**, 301–312.
